# Empagliflozin preserves cardiac function and modulates metabolism in a mouse model of Duchenne muscular dystrophy

**DOI:** 10.64898/2026.03.11.710889

**Authors:** Benjamin J. Zeidler, Connor Thomas, John P. Salvas, Areli Jannes S. Javier, Alyssa M. Richards, Linda A. Bean, Conner C. Earl, Akanksha Agrawal, Niharika Narra, Lifan Zeng, Carol A. Witczak, Joshua R. Huot, Il-Man Kim, Meena S. Madhur, Mark C. Kowala, Larry W. Markham, Craig J. Goergen, Steven S. Welc

## Abstract

Duchenne muscular dystrophy (DMD) is a fatal genetic disorder characterized by skeletal muscle degeneration and cardiomyopathy without a cure. This study examined the therapeutic potential of the sodium-glucose cotransporter 2 (SGLT2) inhibitor empagliflozin (EMPA) on cardiac function in the dystrophin-deficient *mdx* mouse model of DMD. Male mice were fed control chow or EMPA-containing chow (∼25 mg/kg/day), and cardiac function was evaluated longitudinally by four-dimensional ultrasound imaging. EMPA did not alter left ventricular mass or chamber volume but preserved ejection fraction (EF) for 12 weeks, maintained significantly higher EF through 24 weeks, and attenuated global impairment of systolic and diastolic myocardial deformation. These functional improvements were accompanied by reduced cardiomyocyte hypertrophy and decreased expression of cardiac stress genes. EMPA reduced mitochondrial DNA damage, increased mitochondrial DNA copy number, and induced transcriptional signatures consistent with enhanced fatty acid and ketone metabolism, contributing to increased myocardial ATP content. Systemically, EMPA improved body mass trajectory, preserved relative lean mass, enhanced skeletal muscle torque, and did not adversely affect renal function. Together, these findings demonstrate that EMPA improves cardiac performance and mitochondrial integrity while enhancing myocardial energy availability in *mdx* mice, supporting SGLT2 inhibitors as a promising therapeutic strategy for individuals with DMD.

## Introduction

Duchenne muscular dystrophy (DMD) is an inherited disorder caused by mutations in the gene encoding dystrophin, a structural protein essential for cardiac and skeletal muscle integrity and function (1). Affecting approximately 1 in 5,000 live male births (2), DMD initially presents with progressive skeletal muscle weakness, including delayed motor milestones, loss of ambulation, and respiratory insufficiency, with clinical cardiomyopathy emerging later in the disease course. Untreated, DMD is fatal in the late teens to early twenties (3). Improvements in clinical management and skeletal muscle-directed therapies have extended lifespan in DMD, unmasking cardiomyopathy as highly penetrant and dominant determinant of life expectancy (4). Although genomic strategies aimed at restoring dystrophin show promise, their effectiveness in preventing or treating cardiomyopathy remains in question (5). Moreover, current pharmacologic therapies target the renin-angiotensin-aldosterone and sympathetic nervous systems and provide only modest benefit (6), leaving cardiomyopathy as the primary driver of morbidity and mortality in DMD.

Despite imaging studies suggesting that cardiomyopathy begins in the first, the delayed onset of progression observed in DMD patients, typically through the second or third decade, suggests that secondary mechanisms downstream of dystrophin-deficiency drive a pathologic cardiac phenotype. At the cellular level, loss of dystrophin destabilizes the sarcolemma, resulting in increased membrane permeability, dysregulated sodium and calcium homeostasis, and enhanced membrane fragility (7–9). Cardiomyopathy progression is marked by calcium overload, profound mitochondrial dysfunction, metabolic derangements, energetic deficits, and fibrosis (10–14). Importantly, investigations in *mdx* mice, which are also dystrophin null mutants, demonstrate that mitochondrial dysfunction and metabolic abnormalities precede overt cardiac remodeling and dysfunction, implicating impaired energetics as an early driver of disease (15, 16).

Sodium–glucose cotransporter 2 inhibitors (SGLT2i), including empagliflozin (EMPA), have emerged as potent cardioprotective agents. Beyond glycemic control, SGLT2i reduce heart failure hospitalizations, composite cardiovascular outcomes, and both cardiovascular and all-cause mortality in diabetic and non-diabetic heart failure populations (17–21). Mechanistically, SGLT2i induce osmotic diuresis and natriuresis, reprogram substrate availability, enhance cardiac energetic efficiency, and reduce intracellular sodium accumulation, improving calcium handling and preserving mitochondrial function (22). Thus, the diverse and pleiotropic effects of SGLT2i converge with core molecular drivers of dystrophinopathy.

This study was designed to test the hypothesis that EMPA is safe and effective for the treatment of dystrophin-deficient cardiomyopathy. Although the safety and tolerability of SGLT2i are being evaluated in humans including the DuCHennE caRdiomyopathy mItigation Sglt2 inHibitor (CHERISH) clinical trial (NCT07172971) and Repurposing Empagliflozin for DMD-associated Cardiomyopathy clinical trials (NCT06643442), *in vivo* evidence demonstrating preservation of dystrophin-deficient cardiac function and support for mechanistic effects in cardiac tissues are needed. To address this gap, *mdx* mice were treated with EMPA for 24 weeks. Left ventricular (LV) systolic function and global and regional cardiac kinematics were serially assessed using high-frequency four-dimensional ultrasound (4DUS), allowing for the reconstruction of cardiac strain maps. Additional studies evaluated effects on cardiac cellular remodeling, transcriptional regulation of stress and metabolic pathways, mitochondrial adaptations, and myocardial ATP. Subsequent analyses examined whether local cardiac effects were associated with changes in systemic metabolic substrates, regulatory hormones, growth, and body composition. The broader therapeutic potential of EMPA in muscular dystrophy was further evaluated by examining its effects on renal and skeletal muscle dysfunction. The results suggest that SGLT2i may be a viable approach for the treatment of DMD-associated cardiomyopathy.

## Results

### Empagliflozin administration, confirmation of pharmacodynamic activity, and growth effects in mdx mice

The effects of EMPA on *mdx* dystrophy were evaluated in a series of experiments, outlined in Fig. 1A. Importantly, no mortality occurred after initiating treatment. After 24 weeks, *mdx* mice receiving ∼25 mg/kg/day EMPA in feed achieved a mean plasma concentration of 9.53 ± 5.45 ng/mL (Fig. 1B), within the reported therapeutic range in mice (23). EMPA was detectable by Liquid Chromatography-Mass Spectrometry in all treated *mdx* mice, with concentrations exceeding the lower limit of detection (1.01 ng/mL). Pharmacodynamic activity was confirmed by the presence of glucosuria (Fig. 1C). Increased urinary glucose excretion was not associated with changes in absolute body mass in *mdx* mice (Fig. 1D). As expected, *mdx* mice exhibited reduced body mass relative to wild-type (WT) mice, reflecting in part diminished skeletal muscle mass in muscular dystrophy. Because *mdx* mice randomly assigned for EMPA treatment had a lower mean body mass at baseline, body mass data was additionally expressed as relative change from baseline (Fig. 1E). Two-way ANOVA revealed significant effects of EMPA [F(1,17) = 6.1, *p* < 0.05], time [F(1.547, 26.3) = 129.9, *p* < 0.0001], and their interaction [F(1.547, 26.3) = 5.87, *p* < 0.05], supporting a potential effect of EMPA on growth trajectories in *mdx* mice.

**Figure 1:**
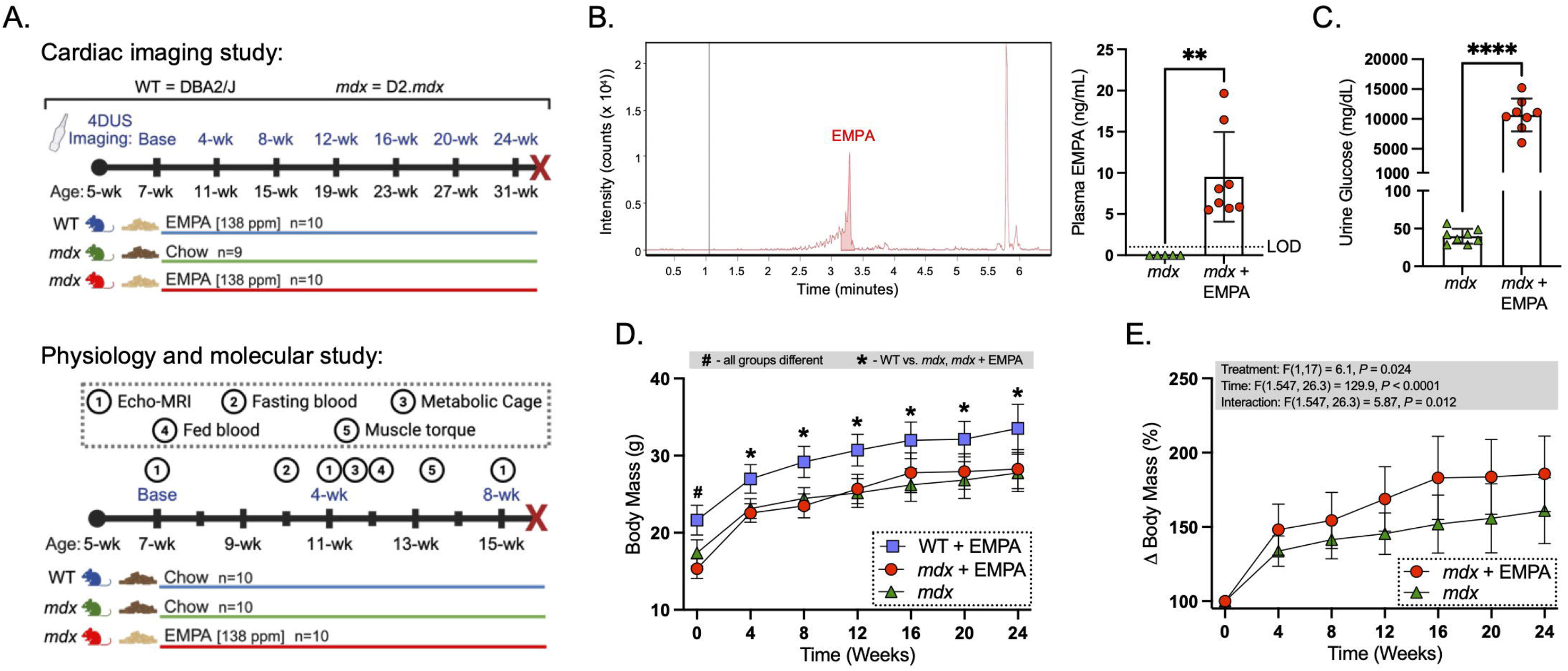
Study design, pharmacodynamic activity and growth effects of empagliflozin in *mdx* mice. (A) Schematic illustrating the long-term empagliflozin (EMPA) treatment protocol with the 4D ultrasound (4DUS) imaging timeline (top), and the short-term EMPA treatment protocol for physiological and molecular characterization (bottom). (B) Representative LC–MS chromatogram of plasma EMPA (left) and quantification of plasma concentrations (right), demonstrating detectable levels in EMPA-treated *mdx* mice (mean 9.5 ng/mL, n = 8) and no detectable levels in untreated *mdx* controls (n = 5; lower limit of detection 1.01 ng/mL). (C) Twenty-four–hour urinary glucose excretion demonstrating pharmacodynamic activity (n = 8 per group). (D) Absolute body mass and (E) relative body mass measured at baseline and every 4 weeks (n = 9–10 per group). Data are presented as mean ± SD. Two-group comparisons were analyzed using two-tailed unpaired t tests. ** *P* < 0.01 and **** *P* < 0.0001. Longitudinal comparisons were analyzed by two-way repeated-measures ANOVA with Tukey’s post hoc test. Statistical significance: *, *P* < 0.05, WT vs. *mdx* and *mdx* + EMPA; #, *P* < 0.05, all groups significantly different from one another.

### Empagliflozin delays the onset of global left ventricular systolic dysfunction and sustains cardiac performance in muscular dystrophy

To evaluate the effects of EMPA on cardiac function in muscular dystrophy, LV function was assessed longitudinally using 4DUS (Fig. 2A). Baseline measurements were obtained prior to treatment at 7 weeks of age, with follow-up assessments conducted every 4 weeks through 31 weeks of age. LV ejection fraction (LVEF) was assessed as a metric of global ventricular systolic function (Fig. 2B). At baseline, LVEF was within the normal range, with no significant difference between groups. LVEF remained stable in EMPA-treated WT mice (61.3-65.3%) throughout the duration of our study, indicating that EMPA did not adversely affect cardiac function in normal, healthy mice. In contrast, *mdx* control mice exhibited a progressive decline in LVEF, decreasing from baseline (65.3 ± 3.3%) to 4 weeks (54.0 ± 3.1%), 8 weeks (48.5 ± 4.5%), 12 weeks (46.4 ± 4.6%), 16 weeks (44.2 ± 6.2%), 20 weeks (41.2 ± 6.3%), and 24 weeks (34.7 ± 4.4%). EMPA delayed the onset of systolic dysfunction in *mdx* mice, preserving LVEF for at least 12 weeks and maintaining significantly higher LVEF thereafter, including at 16 weeks (53.4 ± 3.9% vs. 44.2 ± 6.2%, *p* < 0.01), 20 weeks (50.0 ± 5.0% vs. 41.2 ± 6.3%, *p* < 0.05), and 24 weeks (50.0 ± 3.9% vs. 34.7 ± 4.4%, *p* < 0.0001) compared with *mdx* mice assigned the control diet. Stroke volume (SV) was also measured to assess systolic function (Fig. 2C). Following baseline measurements, SV remained lower in *mdx* controls compared to EMPA-treated WT mice. However, SV was consistently higher in *mdx* mice treated with EMPA compared to control at 4 weeks (28.3 ± 5.9 µl vs. 21.8 ± 4.4 µl, *p* < 0.05), 12 weeks (25.2 ± 6.5 µl vs. 19.7 ± 5.9 µl, *p* < 0.01), 20 weeks (25.3 ± 7.1 µl vs. 17.0 ± 3.9 µl, *p* < 0.05), and 24 weeks (26.3 ± 5.0 µl vs. 18.6 ± 4.0 µl, *p* < 0.01). These differences in SV occurred independently of consistent changes in normalized LV mass (Fig. 2D), heart mass and differences in LV volume (Supplementary Fig. 1).

**Figure 2:**
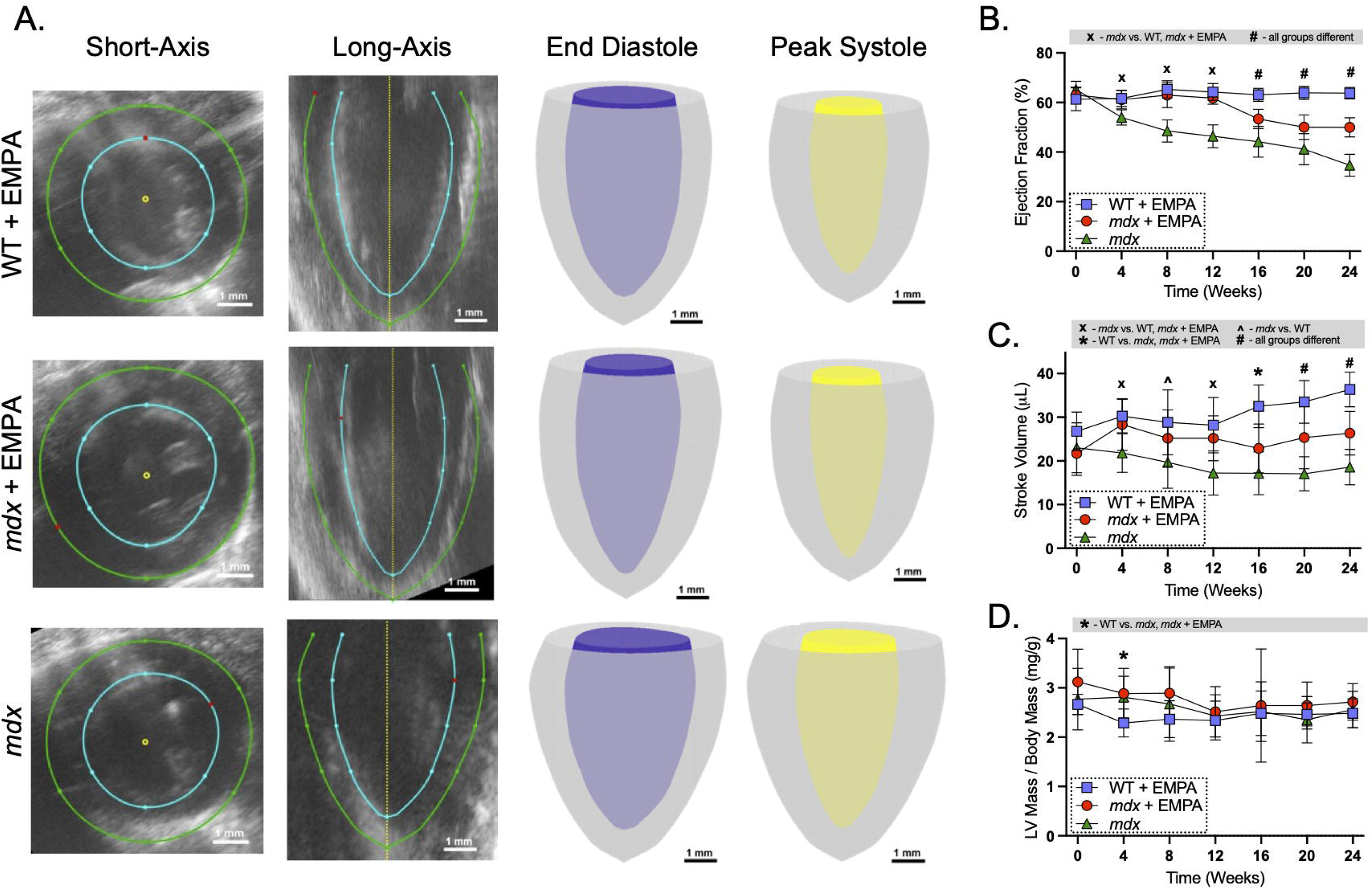
Empagliflozin preserves global left ventricular systolic function in *mdx* mice. (A) Representative week 16 4DUS images generated using custom analysis software. Example segmentations of short-axis and apical long-axis views of the left ventricle (LV), and reconstructed 3D LV models at end-diastole and peak systole. (B) Longitudinal ejection fraction (EF) over 24 weeks. (C) Longitudinal stroke volume over 24 weeks. (D) LV mass normalized to body mass over 24 weeks. Data are presented as mean ± SD. Longitudinal comparisons were analyzed using two-way repeated-measures ANOVA with Tukey’s post hoc test. Statistical significance: x, *P* < 0.05, *mdx* vs. WT and *mdx* + EMPA; ^, *P* < 0.05, *mdx* vs. WT; *, *P* < 0.05, WT vs. *mdx* and *mdx* + EMPA; #, *P* < 0.05, all groups significantly different from one another.

### Empagliflozin preserves myocardial deformation in mdx mice

Detailed myocardial deformation analysis was performed throughout the treatment period using an established 4DUS technique (24), enabling evaluation of global and regional myocardial mechanics. Longitudinal strain trajectories, which depict regional myocardial shortening along the long axis across the cardiac cycle, are shown in Fig. 3A. A visually discernible decline in peak longitudinal strain was evident in *mdx* mice by 4 weeks, whereas a comparable decline in *mdx* mice treated with EMPA was not observed until 16 weeks. In contrast, longitudinal strain trajectories remained relatively stable in EMPA-treated WT mice throughout the study. Strain maps depicting differences in myocardial strain between groups are presented in Supplemental Fig. 2.

**Figure 3:**
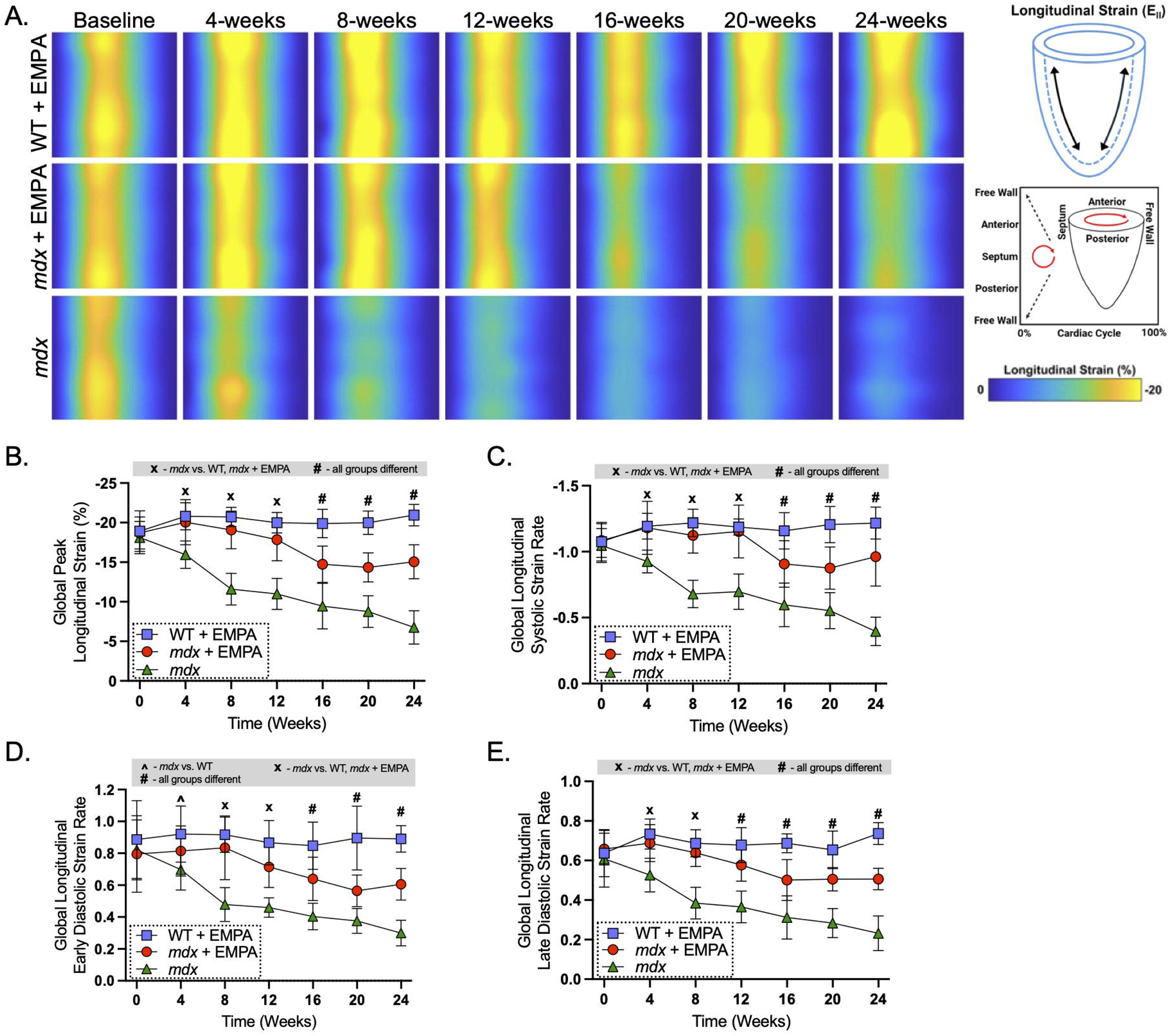
Empagliflozin sustains global longitudinal strain and strain rate in *mdx* mice. (A) Group-averaged heatmaps of LV longitudinal strain at each time point, depicting regional strain patterns (y-axis: free wall, anterior, septum, posterior, and free wall) across one cardiac cycle (x-axis). Color scale denotes low (blue) and high (yellow) longitudinal strain. (B) Global peak longitudinal strain over time. (C) Global systolic longitudinal strain rate. (D) Global early diastolic longitudinal strain rate. (E) Global late diastolic longitudinal strain rate. Data are presented as mean ± SD. N = 9-10 per group. Multi-group comparisons were analyzed by two-way repeated-measures ANOVA with Tukey’s post hoc test for multiple comparisons. Statistical significance: x, *P* < 0.05, *mdx* vs. WT and *mdx* + EMPA; ^, *P* < 0.05, *mdx* vs. WT; #, *P* < 0.05, all groups significantly different from one another.

To assess magnitude and kinetics of myocardial deformation, global peak strain and global longitudinal systolic, early diastolic, and late diastolic strain rates were quantified. At baseline, peak longitudinal strain and associated strain rates were comparable across groups and remained stable in WT mice treated with EMPA throughout the 24 weeklong study (Fig. 3B-E). In *mdx* mice, peak longitudinal strain significantly worsened from baseline values (-18.1 ± 2.0%) to 8 weeks (-11.6 ± 2.0%, *p* < 0.01) and progressively deteriorated through 24 weeks (-6.7 ± 2.1%, *p* < 0.0001). Systolic, early diastolic, and late diastolic longitudinal strain rates exhibited a similar pattern of progressive decline in *mdx* mice over the course of the study (Fig. 3B-E). Overall, these findings indicate global impairment of myocardial deformation during both systole and diastole in *mdx* mice.

EMPA preserved peak longitudinal strain in *mdx* mice at levels comparable to WT mice through at least 12 weeks (Fig. 3B). Although a partial decline occurred thereafter, peak strain remained significantly greater than in *mdx* control mice throughout the remainder of the study. Similarly, EMPA-treated *mdx* mice exhibited preserved systolic and late diastolic strain rates through 12 weeks, after which these measures began to diverge from WT levels but remained significantly greater than *mdx* controls for the rest of the treatment period. Early diastolic strain rate showed a similar pattern, diverging at 8 weeks while remaining preserved through study completion (Fig. 3C-E).

Because cardiac pathology in muscular dystrophy is spatially heterogeneous and characterized by focal pathologies, regional analysis of myocardial deformation was performed to fully capture disease and treatment-related changes. Peak strain and strain rate metrics were segmented across six LV regions: anterior wall, anterior free wall, posterior free wall, posterior wall, posterior septum, and anterior septum (Supplementary Fig. 3). Regional longitudinal peak strain, systolic and diastolic strain rates exhibited temporal patterns that paralleled global deformation metrics, suggesting a homogeneous effect of both dystrophin-deficiency and EMPA treatment on cardiac strain throughout the LV in this study.

To provide a comprehensive assessment of myocardial deformation, circumferential and surface area strains were also quantified (Supplementary Fig. 4), and circumferential strain metrics were further analyzed by region (Supplementary Fig. 5). Peak circumferential and surface area strain, along with corresponding systolic and diastolic strain rates, remained stable in EMPA-treated WT mice but progressively declined in *mdx* mice over the course of the study. EMPA treatment in *mdx* mice preserved circumferential and surface area peak strain and strain rates at levels comparable to the WT group through 12 weeks and remained significantly greater than those observed in *mdx* controls through 24 weeks. The similar patterns observed across longitudinal, circumferential, and surface area strain measures indicate that myocardial deformation abnormalities and treatment effects were consistent across contractile orientations. Collectively, these findings demonstrate that EMPA attenuates global impairments in myocardial deformation in *mdx* mice by preserving systolic shortening and improving both active relaxation and late diastolic filling dynamics.

### Effects of empagliflozin on dystrophin-deficient cardiac remodeling

In the absence of prominent LV chamber remodeling over the 24-week treatment period (Supplementary Fig. 1), cellular- and tissue-level markers of remodeling were assessed, focusing on changes occurring over the initial 8 weeks of treatment (experiments outlined in Fig. 1A), when LVEF remained fully preserved in EMPA-treated *mdx* mice (Fig. 2B). Mean cardiac myocyte cross-sectional area was increased in *mdx* mice compared to WT controls (413.8 ± 70.6 µm^2^ vs. 333.0 ± 21.8 µm^2^, *p* < 0.05) and was significantly reduced by EMPA treatment in *mdx* mice (311.1 ± 34.2 µm^2^, *p* < 0.05) (Fig. 4A). Myocardial fibrosis, assessed by picrosirius red staining, was minimal at this early disease stage, and no differences in collagen content were detected with EMPA treatment in *mdx* mice (Fig. 4B). Collectively, these findings support an association between EMPA-mediated preservation of LV function and attenuation of cardiomyocyte hypertrophy in the absence of overt fibrotic remodeling in *mdx* mice.

**Figure 4:**
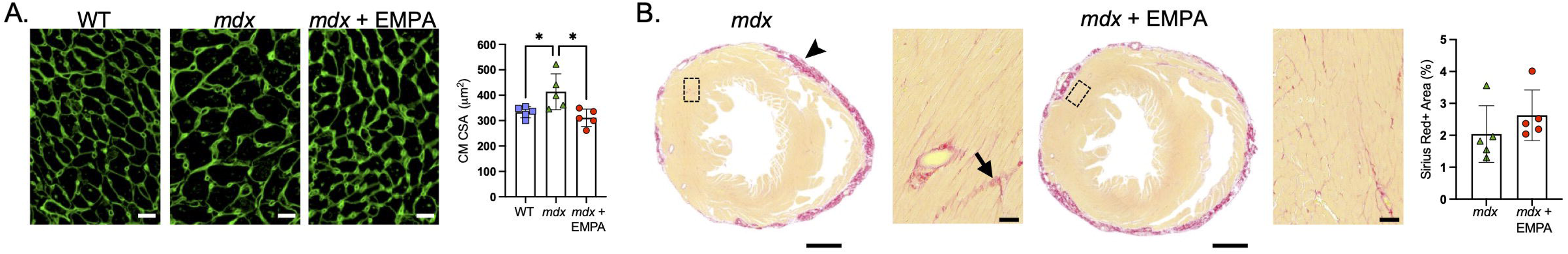
Effect of empagliflozin on pathological cardiac remodeling in *mdx* mice. (A) Representative myocardial sections stained with wheat germ agglutinin (WGA) and quantification of cardiomyocyte (CM) cross-sectional area (CSA). Scale bar: 20 μm. (B) Representative mid-chamber ventricular sections stained with picrosirius red with corresponding quantification of interstitial collagen volume fraction. Scale bars: 1 mm (montage) and 50 μm (insets). Arrowhead indicates collagen associated with spontaneous epicardial calcinosis typical of mice on the DBA/2 background; arrow indicates interstitial myocardial fibrosis. Data are presented as mean ± SD. N = 5 per group. Multi-group comparisons were analyzed by ordinary one-way ANOVA with Tukey’s post hoc test for multiple comparisons or two-group comparisons were analyzed using two-tailed unpaired *t* tests where appropriate. * *P* < 0.05.

### Empagliflozin modulates molecular markers of cardiac stress, fibrosis and metabolism

Early molecular adaptations during the first 8 weeks of EMPA treatment were examined in relation to preserved LV function in *mdx* mice. To determine whether attenuation of cardiomyocyte hypertrophy and maintenance of cardiac performance were accompanied by changes in myocardial stress signaling, qPCR assays were performed to assess transcriptional expression of cardiac stress genes *Nppa*, *Nppb*, and *Myh7.* Myocardial *Nppa* mRNA expression was increased by 30% in *mdx* hearts compared with WT controls and was normalized by EMPA treatment (Fig. 5A). In contrast, *Nppb* mRNA expression was reduced in *mdx* hearts relative to WT mice and was not affected by EMPA. Expression of *Myh6* and *Myh7* transcripts, encoding adult and fetal myosin heavy chain isoforms, was increased by 52% and 97% in *mdx* mice relative to WT controls, respectively. EMPA treatment reduced *Myh7* mRNA expression by 16%. Despite normalization of *Nppa* transcripts and partial reduction of *Myh7* expression, the elevated Myh7/Myh6 transcript ratio in *mdx* hearts remained unchanged by EMPA, indicating attenuation of the cardiac stress response without reversal of the fetal contractile gene program.

**Figure 5:**
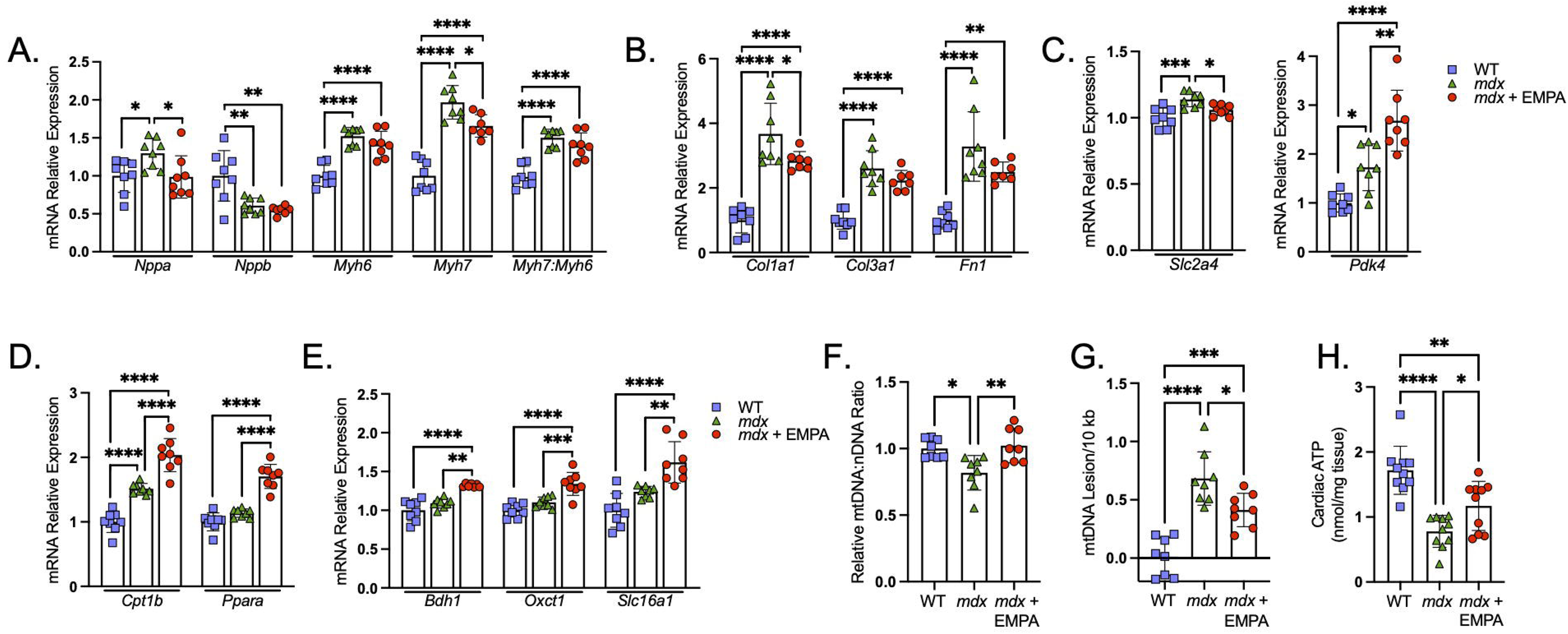
Empagliflozin modulates myocardial remodeling and metabolic gene expression, mitochondrial integrity, and ATP content in *mdx* mice. Myocardial mRNA expression of markers of (A) cardiac stress, (B) fibrosis, (C) glucose metabolism, (D) fatty acid metabolism, and (E) ketone metabolism was quantified by qRT-PCR. (F) Mitochondrial DNA content assessed as the ratio of mitochondrial 16S rRNA to nuclear Hk2 quantified by qRT-PCR. (G) Mitochondrial DNA damage assessed by long-amplicon quantitative PCR. (H) Cardiac ATP content normalized to tissue mass. Data are presented as mean ± SD. N = 7-8 per group. Multi-group comparisons were performed using ordinary one-way ANOVA with Tukey’s post hoc test. * *P* < 0.05, ** *P* < 0.01, *** *P* < 0.001, **** *P* < 0.0001.

Although interstitial myocardial fibrosis was minimal at this stage of disease (Fig. 4B), expression of genes encoding core extracellular matrix components was increased in *mdx* hearts, with *Col1a1*, *Col3a1*, and *Fn1* mRNA levels elevated 3.7-, 2.6-, and 3.3-fold, respectively. EMPA treatment significantly reduced *Col1a1* mRNA expression by 21% (Fig. 5B), consistent with attenuation of early fibrotic gene expression.

Because alterations in mitochondrial function and energy substrate metabolism precede overt cardiac dysfunction and tissue remodeling in dystrophin-deficient mice (15, 16), transcriptional markers of altered cardiac substrate utilization were assessed. Consistent with altered regulation of glucose transport and oxidation, EMPA reduced glucose transporter 4 (*Slc2a4*) mRNA expression and increased pyruvate dehydrogenase kinase 4 (*Pdk4*) expression in *mdx* mice (Fig. 5C). PDK4 is an inhibitor of pyruvate dehydrogenase which regulates entry of carbohydrate intermediates into the Krebs cycle. EMPA also increased expression of genes involved in alternative oxidative substrate utilization, including carnitine palmitoyltransferase 1β (*Cpt1b*) and peroxisome proliferator-activated receptor α (*Ppara*), key regulators of mitochondrial fatty acid transport and β-oxidation (Fig. 5D). In addition, expression of β-hydroxybutyrate dehydrogenase (*Bdh1*), succinyl-CoA:3-ketoacid CoA transferase (*Oxct1*), and the ketone transporter monocarboxylate transporter 1 (*Slc16a1*), consistent with enhanced ketone body uptake and oxidation (Fig. 5E). Gene expression in *mdx* compared to WT hearts did not demonstrate a coordinated transcriptional shift toward reduced fatty acid oxidation, consistent with prior reports of altered substrate utilization in the absence of a clear transcriptional signature (25).

To determine whether transcriptional changes were associated with mitochondrial adaptations, mitochondrial DNA (mtDNA) copy number, mtDNA damage, and myocardial ATP content were evaluated. The mtDNA to nuclear DNA (mtDNA/nDNA) ratio was reduced and mtDNA damage was increased in *mdx* hearts. EMPA treatment normalized mtDNA/nDNA ratio (Fig. 5F) and attenuated mtDNA damage (Fig. 5G). Collectively, these metabolic and mitochondrial adaptations were associated with increased myocardial ATP content (Fig. 5H), indicating improved mitochondrial integrity and enhanced energy availability with EMPA treatment in *mdx* mice.

### Empagliflozin affects body composition and systemic metabolism in muscular dystrophy

SGLT2 inhibition promotes urinary glucose excretion and induces adaptive changes in whole body metabolism, including alterations in glucose homeostasis, hormonal signaling, substrate utilization, and energy expenditure (22). Given the effects of EMPA on cardiac metabolic gene expression and myocardial ATP, systemic metabolic responses to treatment were evaluated to determine whether these cardiac-specific adaptations were accompanied by alterations in whole body metabolic state.

Body mass and composition were evaluated longitudinally at baseline and after 4 and 8 weeks of treatment. Body mass was lower in *mdx* mice compared with WT controls, reflecting marked reductions in both absolute lean mass and fat mass (Fig. 6A). Although body mass was unchanged with short-term EMPA treatment, significant alterations in body composition were observed. EMPA reduced absolute fat mass in *mdx* mice within 4 weeks, resulting in a relative increase in percent lean mass and a corresponding decrease in percent fat mass (Fig. 6A).

**Figure 6:**
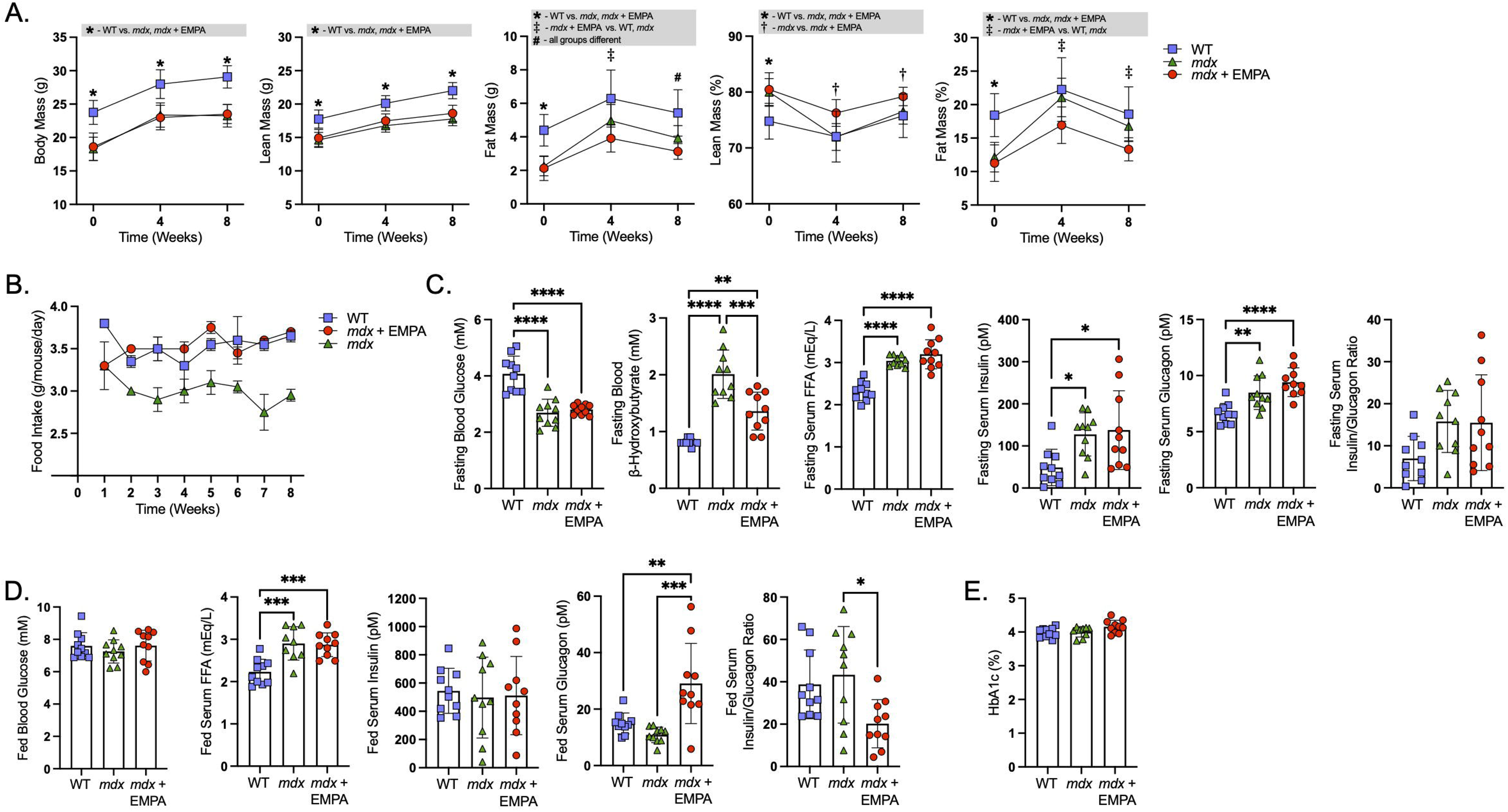
Empagliflozin alters body composition and systemic metabolic parameters in *mdx* mice. (A) Body mass and composition assessed by EchoMRI. Data are presented as mean ± SD. N = 9-10 per group. Multi-group comparisons were analyzed by two-way repeated-measures ANOVA with Tukey’s post hoc test. Statistical significance: *, *P* < 0.05, WT vs. *mdx* and *mdx* + EMPA; †, *P* < 0.05, *mdx* vs. *mdx* + EMPA; ‡, *P* < 0.05, *mdx* + EMPA vs. WT and *mdx*; #, *P* < 0.05, all groups significantly different. (B) Average daily food intake per mouse. Blood glucose, β-hydroxybutyrate, free fatty acids (FFA), insulin, glucagon, and insulin-to-glucagon ratio measured under (C) overnight-fasted and (D) ad libitum–fed conditions. β-hydroxybutyrate values under fed conditions were at the lower limit of detection (0.2 mM). (E) Glycated hemoglobin (HbA1c) was measured after 8 weeks of treatment as an index of chronic glycemia. Data are presented as mean ± SD. N = 9-10 per group. Multi-group comparisons were performed using ordinary one-way ANOVA with Tukey’s post hoc test. * *P* < 0.05, ** *P* < 0.01, *** *P* < 0.001, **** *P* < 0.0001.

Mice were group-housed, and weekly food consumption was normalized to daily intake per mouse. Average daily food intake was lower in *mdx* mice compared to WT controls and was normalized to WT levels with EMPA treatment (Fig. 6B). Together, these findings indicate a selective reduction in fat mass with a relative preservation of lean mass, accompanied by increased food intake that may compensate for caloric loss due to glucosuria.

Systemic metabolites and regulatory hormones were assessed under fasting and fed conditions. After an overnight fast, *mdx* mice exhibited a distinct metabolic profile compared with WT controls (Fig. 6C), characterized by lower blood glucose levels (2.7 ± 0.5 mM vs. 4.1 ± 0.6 mM, *p* < 0.0001), elevated β-hydroxybutyrate (2.0 ± 0.4 mM vs. 0.7 ± 0.1 mM, *p* < 0.0001), and increased circulating free fatty acids (FFAs) (3.0 ± 0.1 mEq/L vs. 2.3 ± 0.2 mEq/L, *p* < 0.0001). These findings are consistent with increased reliance on lipid and ketone-derived substrates in fasting conditions. Fasting insulin (127.5 ± 51.2 pM vs. 48.9 ± 51.2 pM, *p* < 0.05) and glucagon (8.5 ± 1.5 pM vs. 6.6 ± 0.9 pM, *p* < 0.01) were observed in *mdx* mice compared with WT controls, whereas the insulin-to-glucagon ratio was unchanged, suggesting elevated hormone levels without a shift in hormonal balance. EMPA treatment modified fasting substrate availability in *mdx* mice, reducing fasting β-hydroxybutyrate levels (Fig. 6C).

In the fed state, blood glucose concentrations did not differ between groups (Fig. 6D), and β-hydroxybutyrate values were at the lower limit of detection across all conditions (data not shown, 0.2 mM all groups). Serum FFAs remained elevated in *mdx* (2.9 ± 0.4 mEq/L, *p* < 0.001) and EMPA-treated *mdx* mice (2.9 ± 0.3 mEq/L, *p* < 0.001) compared to WT controls (2.2 ± 0.3 mEq/L). While insulin and glucagon levels were similar between WT and *mdx* mice, EMPA increased serum glucagon levels (29.1 ± 14.2 pM vs. 10.9 ± 14.2 pM, *p* < 0.05) and lowered insulin-to-glucagon ratio (*p* < 0.05) in *mdx* mice, indicating a relative shift toward glucagon-dominant signaling. HbA1c levels were unchanged across groups (Fig. 6E), indicating no effects of *mdx* dystrophy or EMPA treatment on long-term glycemic control.

Collectively, EMPA treatment was associated with changes in body composition, substrate availability, and hormonal signaling in *mdx* mice without altering chronic glycemia. These systemic adaptations, together with transcriptional signatures of altered myocardial substrate utilization, suggest alterations in metabolic pathways that may be associated with improved cardiac energy availability and preserved LV function in muscular dystrophy.

### Safety and tolerability of empagliflozin in mdx mice

EMPA acts directly on the kidney, where SGLT2 is abundantly expressed, and promotes osmotic diuresis, which can alter fluid balance and renal tubular function (26, 27). In addition, chronic muscle damage associated with dystrophin loss may secondarily affect renal function (28, 29). To evaluate renal and fluid balance effects of EMPA in muscular dystrophy, water intake, urine output, electrolyte excretion, and markers of renal function and injury were assessed.

After 4 weeks of EMPA treatment, mice were single housed in metabolic cages for 24-hour collections. Urine output did not differ between WT and *mdx* mice. As expected, EMPA induced diuresis, resulting in an approximate 1.4-fold increase in urine output in *mdx* mice (Fig. 7A). Diuresis was at least partially compensated for by an increase in fluid intake (Fig. 7B). *Mdx* control mice consumed 27% less fluid than WT mice in absolute terms, however water intake normalized to body mass was similar between the two groups.

**Figure 7:**
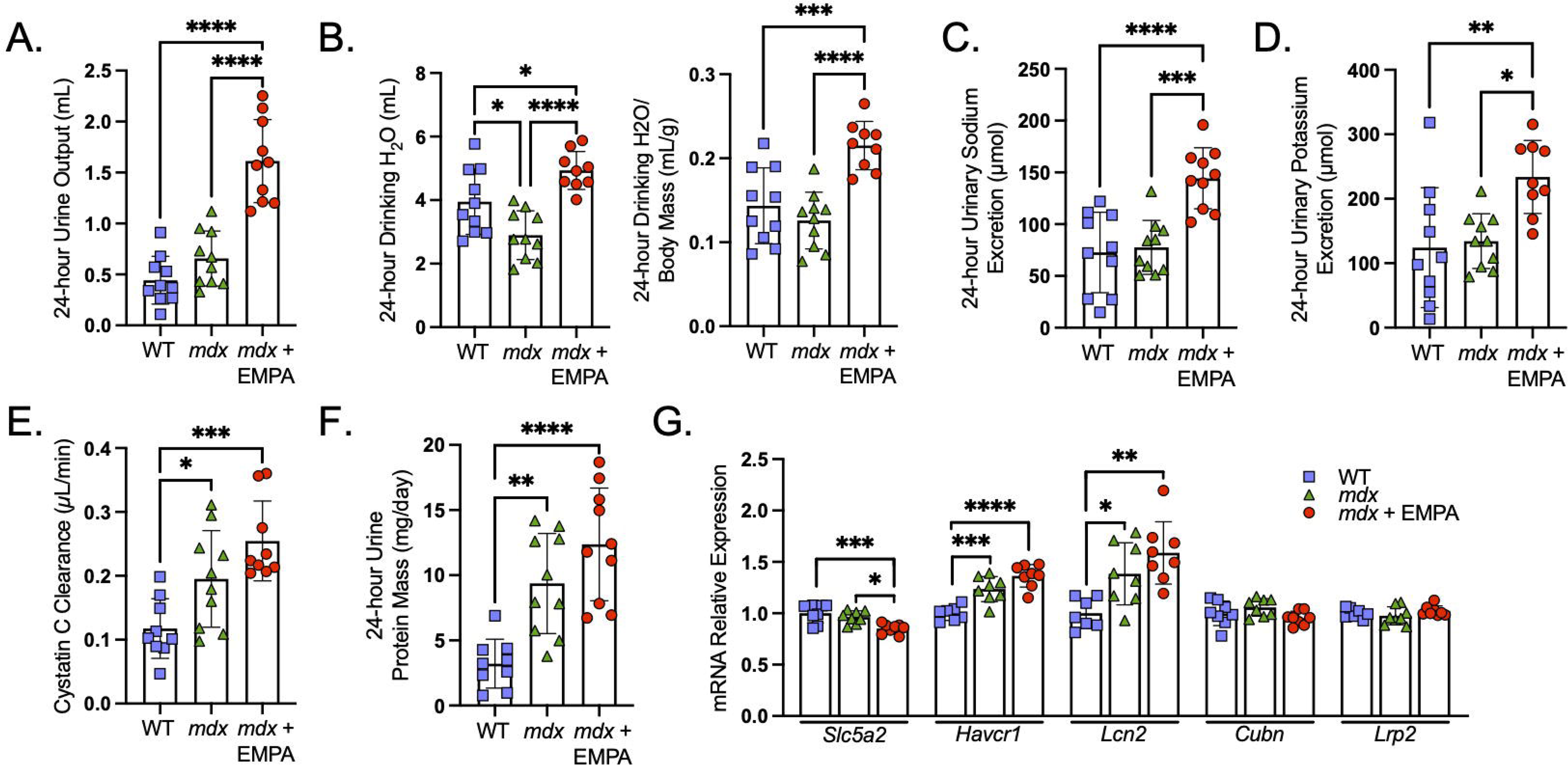
Renal and urinary responses to empagliflozin in *mdx* mice. WT, *mdx*, and EMPA treated *mdx* mice (*mdx* + EMPA) were individually housed in metabolic cages for 24 hours after approximately 5 weeks of treatment. (A) Urine volume. (B) Water intake shown as absolute intake (left) and normalized to body mass (right). (C) Urinary sodium excretion. (D) Urinary potassium excretion. (E) Cystatin C clearance. (F) Total urinary protein excretion. N = 9-10 per group. (G) Renal mRNA expression of SGLT2 (*Slc5a2*) and kidney injury markers by qRT-PCR. N = 7-8 per group. Data are presented as mean ± SD. Multi-group comparisons were performed using ordinary one-way ANOVA with Tukey’s post hoc test. * *P* < 0.05, ** *P* < 0.01, *** *P* < 0.001, **** *P* < 0.0001.

To test if changes in urine volume were accompanied by alterations in electrolyte handling, 24-hour urinary sodium and potassium excretion were measured. Sodium and potassium excretion did not differ between untreated *mdx* and WT mice, suggesting no disruption of tubular electrolyte regulation in *mdx* mice. In contrast, EMPA treatment increased sodium excretion by 86% (Fig. 7C) and potassium excretion by 74% (Fig. 7D) compared with *mdx* controls, consistent with natriuresis and kaliuresis secondary to SGLT2 inhibition.

Because of the marked reductions in muscle mass in *mdx* mice, renal function was estimated by cystatin C clearance rather than creatinine-based measures (30). Cystatin C clearance was elevated in *mdx* mice than in WT controls (Fig. 7E), consistent with increased glomerular filtration. EMPA treatment did not further alter cystatin C clearance, indicating no additional effect on renal filtration in *mdx* mice. Similarly, 24-hour urinary protein excretion was elevated in *mdx* mice relative to WT and was unchanged by EMPA treatment (Fig. 7F). These findings suggest that *mdx* mice exhibit increased filtration and proteinuria consistent with altered renal function associated with muscular dystrophy, but these changes were not exacerbated by EMPA treatment.

Renal safety was further tested by gene expression analyses to assess transcripts associated with proximal tubular function and injury. EMPA treatment reduced *Slc5a2* (SGLT2) mRNAs in *mdx* mice, potentially reflecting a compensatory downregulation of the drug target in response to sustained SGLT2 inhibition (Fig. 7G). Expression of proximal tubular injury markers *Havcr1* (KIM-1) and *Lcn2* (NGAL) was elevated in *mdx* mice compared with WT controls but was not further altered by EMPA treatment. Transcript levels of *Cubn* and *Lrp2*, which mediate proximal tubular protein reabsorption, did not differ between groups suggesting that baseline proteinuria is not associated with transcriptional regulation of proximal tubular reabsorption pathways.

In summary, these findings demonstrate that EMPA induces osmotic diuresis and natriuresis in *mdx* mice consistent with the well-described physiological effects of SGLT2i. Additionally, EMPA treatment did not worsen renal dysfunction or transcriptional markers of tubular injury, supporting its safety and tolerability for use in muscular dystrophy.

### Empagliflozin improves skeletal muscle contractile function in muscular dystrophy

Because DMD is characterized by progressive skeletal muscle degeneration, therapies that target cardiomyopathy must also be evaluated for their effects on skeletal muscle to ensure overall therapeutic benefit. Having established that EMPA delays the onset of LV systolic dysfunction, preserves long-term cardiac performance, and is well tolerated, its impact on skeletal muscle function was next assessed in *mdx* mice.

Serum creatine kinase, a biomarker for muscle injury, was evaluated and found to be elevated in *mdx* mice compared with WT controls but was not further affected by EMPA (Fig. 8A), indicating that EMPA did not exacerbate muscle injury. Given that EMPA alters systemic substrate availability and hormonal signaling, *in vivo* plantar flexion was measured to capture integrated metabolic, neural and vascular influences on skeletal muscle function. Twitch force scaled to body mass was reduced by 50% in *mdx* mice and 38% in EMPA-treated *mdx* mice relative to WT controls (Fig. 8B). Maximum torque normalized to body mass was reduced by 39% in *mdx* mice compared with WT controls. However, EMPA treatment increased maximum torque by 17% relative to *mdx* controls, partially restoring muscle function (Fig. 8C). Reductions in muscle torque in *mdx* mice were associated with decreased tibialis anterior (TA) and extensor digitorum longus (EDL) muscle mass normalized to body mass relative to WT controls. EMPA treatment did not significantly alter normalized TA, EDL or soleus muscle mass compared to *mdx* controls (Fig. 8D). Overall, EMPA treatment partially restored contractile function in *mdx* mice without affecting muscle injury or increasing muscle mass, consistent with improved muscle quality rather than preservation of muscle size.

**Figure 8:**
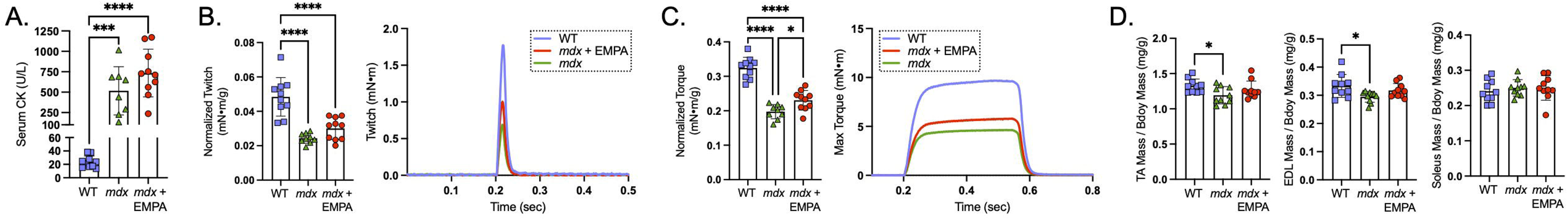
Empagliflozin improves maximal *in vivo* skeletal muscle torque without altering muscle injury or mass. (A) Serum creatine kinase (CK) levels. *In vivo* plantar flexion torque of the triceps surae muscle group was assessed in WT, *mdx*, and *mdx* + EMPA mice after approximately 6 weeks of treatment. (B) Maximal twitch torque normalized to body weight (mN·m/g) with representative twitch traces. (C) Maximal tetanic torque normalized to body weight (mN·m/g) with representative tetanic traces. (D) Muscle mass normalized to body mass of fast-twitch tibialis anterior (TA) and extensor digitorum longus (EDL), and slow-twitch soleus muscles. Data are presented as mean ± SD. Multi-group comparisons were analyzed using ordinary one-way ANOVA with Tukey’s post hoc test. \**P* < 0.05, \*\*\**P* < 0.001, \*\*\*\**P* < 0.0001.

## Discussion

The principal finding of this study is that EMPA delays cardiac functional decline and preserves ventricular performance in dystrophin-deficient mice. This conclusion is supported by longitudinal 4DUS imaging demonstrating preservation of global systolic function, myocardial deformation, and both systolic and diastolic mechanics over an extended disease course. The current results also offer mechanistic insight into how EMPA preserves cardiac function in muscular dystrophy. Functional benefits were accompanied by reduced cardiomyocyte hypertrophy and lower expression of cardiac stress markers, with early improvement preceding overt fibrosis. EMPA treatment was associated with increased myocardial mtDNA copy number, reduced mtDNA damage, transcriptional changes consistent with enhanced fatty acid and ketone oxidation and increased myocardial ATP content. Together, these findings are consistent with improved mitochondrial integrity and myocardial energetic capacity in dystrophin-deficient hearts. In addition, EMPA demonstrated broader therapeutic potential in muscular dystrophy, as reflected by favorable effects on long-term body mass trajectory, preservation of relative lean mass, and enhanced *in vivo* skeletal muscle torque, without adverse effects on renal function in *mdx* mice.

The *in vivo* effects of SGLT2i therapy have not previously been evaluated in dystrophin-deficient cardiomyopathy. However, prior work demonstrated that EMPA treatment restored peak inward sodium current during membrane depolarization by pharmacochaperoning Nav1.5, thereby rescuing its plasma membrane expression in cardiomyocytes and Purkinje fibers derived from *mdx* mice and dystrophin-deficient rats. (31, 32). Reduced peak sodium current in ventricular cardiomyocytes and Purkinje fibers may impair ventricular conduction and pre-dispose to arrhythmias potentially contributing to mortality in DMD (33–35). Whether SGLT2 inhibitors reduce arrhythmia burden, and how such potential electrophysiological effects relate to improvements in LV function observed in the present study, remains to be determined.

EMPA treatment in *mdx* mice promoted a shift toward enhanced adipocyte lipid mobilization and utilization, characterized by preferential reduction in fat mass with preservation of total body mass and increased relative lean mass. These changes were accompanied by a hormonal shift toward glucagon-dominant signaling. Despite a reduced insulin-to-glucagon ratio, fasting β-hydroxybutyrate levels were lower, which may reflect increased peripheral ketone utilization. Increased expression of *Slc16a1*, *Bdh1*, and *Oxct1* in hearts from *mdx* mice treated with EMPA supports enhanced myocardial ketone oxidation capacity. In contrast to prior reports of increased glucose reliance in *mdx* hearts (25), EMPA increased *Cpt1b* and *Ppara* expression and reduced *Slc2a4* and *Pdk4* expression, suggesting a shift toward a more oxidative metabolic phenotype. Thus, EMPA may promote a coordinated shift in systemic substrate availability and myocardial transcriptional reprogramming toward a more oxidative metabolic phenotype that may be advantageous in dystrophic hearts.

Calcium dysregulation, mitochondrial dysfunction, and excessive reactive oxygen species (ROS) generation are all central features of dystrophin-deficient cardiomyopathy. The observed increase in mtDNA copy number and reduction in mtDNA lesion frequency following EMPA treatment are therefore consistent with improved mitochondrial integrity. Enhanced ketone and fatty acid oxidation has been proposed as a mechanism that improves mitochondrial efficiency and reduces electron leak from the respiratory chain, potentially lowering ROS production (36). Additionally, ketone metabolism has been associated with improved redox balance and activation of cytoprotective signaling pathways that may further limit oxidative injury (37). Alternatively, SGLT2i may reduce intracellular sodium accumulation through natriuretic effects or modulation of sodium transport and improve calcium handling, thereby supporting mitochondrial function, lowering metabolic stress, and enhancing contractility (38–42). These mechanisms may collectively contribute to preserved mitochondrial integrity and cardiac performance.

The beneficial effects of EMPA on dystrophic cardiac function may also reflect coordinated cardiorenal protective mechanisms. Individuals with DMD are at increased risk of renal impairment, which can further exacerbate cardiac dysfunction (43). Although EMPA was shown to slow the progression of kidney disease in type 2 diabetes (44), it did not significantly alter Cystatin C clearance, 24-hour urinary total protein excretion, or expression of proximal tubular injury genes (KIM1 and NGAL), suggesting no measurable change in eGFR or injury in this model. Nonetheless, EMPA induced osmotic diuresis and natriuresis. SGLT2i preferentially reduce tissue sodium and interstitial fluid volume, producing a greater reduction in interstitial congestion relative to intravascular volume compared with conventional diuretics (27, 45, 46). Given that sodium accumulation and interstitial fluid expansion may precede cardiac dysfunction in DMD (9, 47), EMPA could alleviate tissue congestion, thereby contributing to improved cardiac performance.

The effects of SGLT2i on skeletal muscle function remain incompletely defined. By promoting glucosuria, SGLT2i induce a negative energy balance that typically results in reductions in body mass and fat mass, raising concerns about potential skeletal muscle wasting (48). However, in the present study, long-term (24-week) EMPA treatment was associated with a relative increase in body mass in *mdx* mice. Experimental data in non-diabetic mice indicate that SGLT2i can reduce muscle mass and function under conditions of caloric restriction (paired feeding), whereas *ad libitum* feeding mitigates these effects, suggesting that muscle loss is secondary to negative energy balance rather than a direct myotoxic effect (49). Consistent with this interpretation, the energetic deficit from glucosuria in our study appeared to be compensated by increased food intake. Short-term (8-week) EMPA treatment did not alter total body mass but resulted in a relative increase in lean mass, accompanied by improved *in vivo* muscle torque without changes in muscle mass, suggesting enhanced muscle quality. Whether these modest benefits reflect systemic metabolic reprogramming or differential effects on skeletal muscle and cardiac muscle remains to be determined.

Loss of dystrophin is associated with systemic metabolic alterations that remain incompletely understood. In the present study, *mdx* mice exhibited enhanced fasting lipolysis and ketogenesis accompanied by lower blood glucose levels. The concurrent elevation of both insulin and glucagon, hormones with opposing metabolic effects, suggest endocrine dysregulation and possible insulin resistance in *mdx* mice. Persistently elevated fed-state FFAs and reduced body and fat mass further imply increased energy expenditure. Consistent with these findings, prior studies have described a hypermetabolic phenotype in *mdx* mice on both C57BL/10 and DBA/2 genetic backgrounds, potentially reflecting the substantial energetic demands of ongoing muscle regeneration compared with WT mice (50, 51). However, correction of skeletal muscle pathology does not rescue the metabolic phenotype, suggesting that metabolic abnormalities may occur independently of skeletal muscle pathology or may be related to dystrophin-deficiency in non-skeletal muscle tissues (51). The mechanisms of this broader metabolic dysregulation downstream of dystrophin-deficiency, as well as the pathways through which EMPA modifies this molecular state, remain to be defined.

To our knowledge, this is among the first studies to apply longitudinal 4DUS imaging to assess LV function in *mdx* mice (24). Compared with conventional two-dimensional methods, four-dimensional imaging captures three-dimensional cardiac structure throughout the cardiac cycle and may be better suited to the heterogeneous myocardial pathologies seen in muscular dystrophy. Using this approach, a progressive decline in LVEF was observed from 7 to 31 weeks of age, indicating an extended period of progressive functional deterioration distinct from previous reports (52). Given that LV dysfunction progresses in patients with DMD, defining this window of decline may enhance the translational relevance of preclinical therapeutic evaluation.

Myocardial strain metrics from four-dimensional imaging data are highly sensitive to both global and regional LV function (53). Globally, circumferential and longitudinal strains follow similar trends across groups, and all regional patterns appear homogeneous in both types of strain, suggesting that strain changes are homogeneous. Previous studies of cardiomyopathy progression in DMD patients using speckle-tracking echocardiography and tissue Doppler imaging, however, have reported more severe changes in longitudinal strain in the basal lateral regions of the LV (54, 55). Additionally, human cardiac magnetic resonance imaging (MRI) with late gadolinium enhancement has exhibited initial onset of fibrosis mainly in the lateral free wall of the LV (56). Reports using cardiac MRI suggest regional heterogeneity in human circumferential strain data, as more pronounced changes were observed in strain and strain rate of the LV base compared to the mid-LV or apex, regardless of DMD status (57, 58). This variation in regional patterns could be due to differences in strain-mapping approaches (ultrasound vs. MRI) or the nature of study participants (mice vs. humans). Notably, we also reported similar regionally homogeneous strain metric patterns in a separate animal study investigating the acceleration of DMD-related cardiomyopathy with isoproterenol, albeit in *mdx* mice on the C57BL10 genetic background (24). One possibility is that while there may be regional strain differences in *mdx* mice, the spatial distribution in the mouse LV may be more heterogenous in individual animals compared to humans, presenting a more consistent trend when averaged across groups. Further investigation into the comparison of regional patterns of cardiac strain in both preclinical and human DMD studies is warranted.

The findings of this study should be interpreted in the context of several limitations. This study characterizes the physiological effects of SGLT2i in muscular dystrophy and the functional impact of EMPA on dystrophin-deficient cardiac function, but this study was not designed to establish causal links between the molecular alterations observed and the improvements in cardiac performance. While multiple molecular changes were detected that may contribute to the observed functional benefits, the upstream mechanisms underlying these effects remain undefined. Studies incorporating genetic models, including SGLT2-deficient mice, would be necessary to determine whether these effects are mediated through on-target or off-target pathways (59). Direct assessment of myocardial substrate utilization was not performed.

Definitive evaluation would require tracer-based metabolic flux analyses, positron emission tomography imaging, and/or mitochondrial respirometry using defined substrates, which were beyond the scope of the present work. EMPA was administered prophylactically and attenuated functional decline over 24 weeks. However, the durability of benefit and the ability to halt or reverse established cardiomyopathy remain unknown. Finally, while the *mdx* mouse model is a genetic homolog of DMD and recapitulates key pathological features, its phenotype is milder than DMD. Accordingly, extrapolation of these findings to clinical populations should be made with caution.

In summary, SGLT2i are being evaluated as potential therapies for DMD-associated cardiomyopathy based on conceptual overlap between DMD pathogenesis and the hypothesized mechanisms of SGLT2 inhibition. The present study provides *in vivo* evidence that EMPA preserves cardiac function and attenuates adverse cellular remodeling in dystrophin-deficient *mdx* mice. Functional benefits were accompanied by increased myocardial ATP content, preservation of mitochondrial integrity, and coordinated metabolic adaptations consistent with improved oxidative capacity. EMPA treatment was also not associated with adverse renal effects and conferred modest improvements in skeletal muscle function. Collectively, these findings support further investigation of EMPA as a potential therapeutic strategy for individuals with DMD.

## Methods

### Sex as a biological variable

Male *mdx* mice were used in this study since DMD is an X-linked recessive disease that predominantly affects males.

### Animals

Two cohorts of 5-week-old DBA/2J (WT; strain #000671) and D2.B10-Dmd*^mdx^*/J (*mdx*; strain #013141) mice were purchased from The Jackson Laboratory (n=60 total). The first cohort of mice were housed at Purdue University for cardiac phenotyping (n=30). One *mdx* mouse died during the acclimation period prior to treatment. At 7 weeks of age, cages of 4-5 *mdx* mice were randomized to receive either chow containing 138 ppm of EMPA (*mdx* + EMPA, n=10) or control chow (*mdx*, n=9) and were maintained on their respective diets until 31 weeks of age (24 weeks of treatment). All WT mice received the EMPA-containing chow (WT + EMPA, n=10). The custom formulated diet (#TD.230471) was produced by Inotiv-Teklad using a standard diet base (#2918) supplemented with EMPA (MedChemExpress, #HY-15409). The diet was formulated based on the observation that *mdx* mice consume approximately 3.1 g/day of feed, yielding a targeted dose of ∼25 mg/kg/day, falling within the previously reported effective dosing range of 10–35 mg/kg/day in mice (23, 60–62). The second cohort of mice was housed at Indiana University School of Medicine and used for metabolic assessments, skeletal muscle function testing, and molecular analyses. At 7 weeks of age, cages of *mdx* mice were randomized to receive EMPA containing chow (*mdx* + EMPA, n=10) or control chow (*mdx*, n=10). WT mice in this cohort received control chow (WT, n=10). Mice in this cohort were treated for 8 weeks, from 7 to 15 weeks of age. Food was stored at 4°C and replaced weekly. All mice had *ad libitum* access to food and water.

### Ultrasound Imaging

At 7 weeks of age, baseline image acquisition was performed, followed by echocardiographic assessments every 4 weeks for 24 weeks. Cardiac imaging was conducted using a Vevo 3100 high-frequency ultrasound system (FUJIFILM VisualSonics) with a 40 MHz center frequency MX550D linear array probe. Mice were anesthetized using 2.0% isoflurane at 0.7-1.0 mL/min and secured on an imaging stage in a supine position. After ventral thorax hair was removed with depilatory cream, internal body temperature was maintained at 37 ± 2°C, and heart rate and respiratory rate were maintained at 400–600 beats per minute and 40–120 breaths per minute, respectively. All vitals were monitored throughout the imaging process and anesthesia delivery, and imaging stage temperature were adjusted to maintain these ranges.

Four-dimensional (4D) echocardiography (3D volume + time) was performed at each imaging timepoint. In the parasternal short axis orientation, we used a linear step motor with step size 0.13mm to scan from the apex up to the ascending aorta at 300 frames/sec, capturing a 3-dimensional video of the LV across one cardiac cycle. Using a custom toolbox in MATLAB 2023a (Mathworks, Inc.), we reoriented each image and labeled the base, apex, and endocardial and epicardial border location across the cardiac cycle. As previously described, the custom toolbox allows for the calculation of cardiac function and morphology metrics (EF, stroke volume, LV mass, etc.) and global and regional myocardial deformation metrics (peak strain, systolic strain rate, early diastolic strain rate, late diastolic strain rate) (24, 63).

### Histopathology

Hearts were fixed in 10% neutral-buffered formalin, processed, and embedded in paraffin. Myocardial fibrosis was assessed in cardiac sections using Picrosirius Red staining, as previously described with modifications (64). Sections were deparaffinized and hydrated, rinsed in distilled water and incubated in decalcification buffer (Formical-2000, StatLab) overnight with gentle shaking. Slides were rinsed with running water for 10 minutes and the tissues were stained in 0.1% Direct Red 80 (Sigma-Aldrich, 365548) prepared in 1.3% picric acid for 1 hour. Following 2 quick acidified water rinses, tissues were dehydrated through graded ethanol, cleared in xylene, and mounted with Cytoseal XYL (Epredia). Cardiomyocyte cross-sectional area was evaluated in mid-ventricular sections stained with FITC-conjugated wheat germ agglutinin (1:100; Vector Laboratories, FL-1021-5). Whole section montages were captured using a Zeiss AxioObserver 7 inverted epifluorescence microscope with motorized stage, ZenBlue 3.3 software, and Zeiss Axiocam 105 (color) and Axiocam 506 (monochrome) cameras. Images were deidentified prior to analysis and quantified in a blinded manner. Fibrosis was quantified by measuring Sirius Red–positive area using ImageJ/FIJI thresholding and expressed as the percentage of total myocardial area. LV cardiomyocyte cross-sectional area was quantified by measuring at least 100 myocytes per section across a minimum of five fields per sample.

### Metabolic Cage

After 4-5 weeks of treatment, mice were individually housed in metabolic cages (Tecniplast, #3600M021) for 24-hour assessment of water intake and urine collection for metabolic analyses.

### Blood sampling and analysis

Blood samples were collected from the tail vein between 8:00–9:00 AM from mice with *ad libitum* access to food or following a 12–14-hour overnight fast, as described previously (65). Blood glucose was measured using a glucometer (OneTouch, Ultra2) and β-hydroxybutyrate levels were measured with a blood ketone meter (KetoBM, #KBM-MK001). Hemoglobin A1c (Roche Diagnostics, Tina-quant HbA1c Gen.3) was measured by the Translational Core Facility at the Indiana University School of Medicine using an automated Cobas Integra 400 Chemistry Analyzer. For serum analyses, blood was allowed to clot at room temperature for 60 minutes and centrifuged at 5,000 × g for 10 minutes. The serum fraction was transferred to low protein-binding microcentrifuge tubes and centrifuged again for 10 minutes to remove residual red blood cells. Serum glucagon (Mercodia, #10-1281-01), insulin (CrystalChem, #90080), and cystatin C (R&D Systems, #MSTC0) were measured by ELISA. Creatine kinase activity (BioAssay Systems, #ECPK-100) and free fatty acids (Fujifilm Wako Pure Chemical Corp., #299-94301) were determined using commercial assay kits.

### Urine analyses

Urinalysis was performed by the Translational Core using a Cobas Integra 400 Chemistry Analyzer. Sodium and potassium were measured by ion-selective electrode methods. Urine glucose was measured using the Glucose HK Gen.3 assay, and total protein was measured using the Total Protein Urine/CSF Gen.3 assay (Roche Diagnostics). Urine cystatin C concentration was measured by ELISA (R&D Systems, #MSTC0). Cystatin C clearance was used for eGFR and was calculated as: cystatin C clearance (mL/min) = (urine cystatin C (mg/L) x urine flow (mL/min)) / (plasma cystatin C (mg/L)).

### Body composition

Mice were weighed to ±0.1 g. Lean and fat mass was assessed using an EchoMRI body composition analyzer, as described previously (66).

### RNA isolation and quantitative Real-Time PCR (qRT-PCR)

RNA isolation and qRT-PCR assays were completed as previously described (64). Briefly, heart and kidney tissues were homogenized in Trizol, and total RNA was extracted with chloroform separation and purified using RNeasy Plus kit with gDNA eliminator columns (Qiagen). RNA was reverse transcribed with Maxima H Minus cDNA Synthesis Master Mix (Thermo Fisher Scientific) on a Bio-Rad C1000 thermal cycler. Primers were designed in-house (Supplementary Table 1) and commercially synthesized (MilliporeSigma). Primer specificity was verified by amplicon size using agarose gel electrophoresis and melt curve analysis for a single peak. Primer efficiency (90–110%) was assessed using serial dilutions of cDNA. qRT-PCR was performed on a Bio-Rad CFX384 real-time system using Sybr Green chemistry. Relative expression of target transcripts was calculated using the geometric mean of two reference genes. *Rps20* and *Psmc4* were used for cardiac samples, and *Ap3d1* and *Ywhaz* were used for kidney samples. Gene expression in WT control samples was normalized to 1, and all other values were expressed relative to controls.

### Total DNA isolation, mitochondrial DNA (mtDNA) damage, and mtDNA to nuclear DNA (nDNA) ratio

Total genomic DNA was isolated from ventricles using Genomic Tips (Qiagen). Purified DNA was quantified fluorometrically using Quanti-iT PicoGreen assay (Invitrogen). Samples were initially diluted to 10 ng/mL and re-quantified, then adjusted to a final concentration of 3 ng/mL. Mitochondrial DNA (mtDNA) damage was assessed by long amplicon quantitative PCR (QPCR) amplification of long (10 kb) and short (117 bp) mtDNA fragments (Supplementary Table 1) using KAPA LongRange HotStart DNA Polymerase (Roche), as described previously (64). Reactions were performed in triplicate, and amplification linearity was verified using a 50% genomic DNA control. PCR products were quantified fluorometrically. Lesion frequency normalized for mitochondrial copy number was calculated as the negative natural log of the relative amplification as performed previously (64). The mtDNA to nDNA ratio was determined by qRT-PCR as previously described (67). The mitochondrial 16S rRNA gene, located within a stable region of the mitochondrial genome and present as a single copy per mtDNA molecule, was used to quantify mtDNA. HK2, a stable and single-copy nuclear gene, served as the normalization control. Relative mtDNA content was calculated by normalizing 16S rRNA amplification to HK2 amplification. Data were expressed relative to the WT control group within each experiment.

### Myocardial ATP

Cardiac tissue samples snap-frozen in liquid nitrogen were used to measure myocardial ATP content using an ATP assay kit (Abcam, ab83355) according to the manufacturer’s instructions. ATP concentrations were normalized to tissue mass for each sample.

### In vivo plantar flexion

Mice underwent *in vivo* plantar flexion torque testing to assess contractility of the triceps surae (Aurora Scientific), as previously described (66). Briefly, mice were anesthetized with 2% isoflurane, the right hindlimb was secured with the foot positioned at 90° to the tibia attached to a force transducer, and the limb was fixed with a knee clamp. Two disposable monopolar electrodes (Natus Neurology) were inserted to stimulate the tibial nerve. Maximal twitch torque was determined using supramaximal 0.2-ms square-wave pulses. Peak tetanic plantar flexion torque was then measured in response to supramaximal stimulation (0.2 ms pulse width) delivered at 125 Hz. Twitch and tetanic torques were analyzed using Dynamic Muscle Control Data Analysis software (v5).

### Liquid Chromatography Mass Spectrometry (LC-MS)

Plasma samples were mixed with HPLC-grade acetonitrile, sonicated for 60 seconds to precipitate proteins, and centrifuged at 4,000 × g for 5 minutes. The supernatant was collected for LC–MS analysis at Indiana University School of Medicine Chemical Genomics Core facility, and 20 µL was injected per run. LC–MS analysis was performed using an Agilent 1290 LC system coupled to an Agilent 6545 QTOF mass spectrometer. Separation was achieved on an Agilent Zorbax Eclipse Plus C18 column (2.1 × 50 mm, 1.8 µm) using 0.1% formic acid in water (mobile phase A) and 0.1% formic acid in acetonitrile (mobile phase B) at a flow rate of 0.6 mL/min. The gradient began at 95% A (5% B) for 0.5 min, increased to 95% B over 4.5 min, held for 0.5 min, returned to 5% B over 0.5 min, and equilibrated for 0.5 min. Mass spectrometry was performed using electrospray ionization in negative ion mode with nitrogen as the nebulizing and drying gas. Instrument settings were: nebulizer pressure 25 psi, drying gas temperature 325°C, drying gas flow 8 L/min, sheath gas temperature 400°C, sheath gas flow 12 L/min, fragmentor 90 V, skimmer 65 V, and Oct 1 RF Vpp 750 V.

EMPA calibration standards were prepared from a 1 mg/mL stock solution in acetonitrile, diluted in 50% acetonitrile to generate six two-fold serial dilutions (6.25–200 pg). Twenty microliters of each standard and blank were injected to generate calibration curves. The limit of detection (LOD) was calculated as:(\bar{S}{blank} + 3.3 \times SD{blank}) / m, where SLJ_blank is the mean blank signal, SD_blank is the standard deviation of the blank, and m is the slope of the calibration curve. The calculated LOD for EMPA was 1.01 ng/mL. EMPA concentrations in plasma samples were quantified using the standard calibration curve.

### Statistics

All data are presented as mean ± standard deviation (SD). All statistical analysis was conducted in Prism 10.2.3 (GraphPad Software). Prior to analysis, statistical outliers were identified using Grubbs’ outlier test. For comparisons between two groups, an unpaired two-tailed Student’s *t*-test was used. For comparisons among three groups, one-way analysis of variance (ANOVA) followed by Tukey’s multiple comparison correction. For longitudinal outcomes, a two-way repeated measures ANOVA was conducted to assess the effects of group, time and their interaction, followed by a Tukey’s multiple comparisons test to evaluate differences between the three experimental groups across time points. Differences with a *p*-value < 0.05 were considered statistically significant. Normality of residuals and equal variance assumptions were verified using the Shapiro-Wilk and Levene’s tests, as well as qualitative inspection of QQ- and residual plots.

### Study Approval

All procedures conducted involving laboratory animals were reviewed and approved by the Institutional Animal Care and Use Committees at Purdue University (protocol 2002002016) and Indiana University School of Medicine (protocol 25052).

## Author contributions

Conceptualization, A.A., L.A.B, C.C.E., C.J.G., A.J.S.J., M.C.K., L.W.M., C.T., S.S.W.; Methodology, L.A.B, C.C.E., C.J.G., J.R.H., A.J.S.J., C.T., A.M.R., J.P.S., C.A.W., B.J.Z., L.Z., S.S.W.; Validation, L.A.B, C.J.G., A.J.S.J., C.T., A.M.R., J.P.S., B.J.Z., L.Z., S.S.W.; Formal Analysis, L.A.B, C.J.G., J.R.H., A.J.S.J., I.K., M.C.K., M.S.M., N.N., C.T., A.M.R., J.P.S., C.A.W., B.J.Z., L.Z., S.S.W.; Investigation, L.A.B, J.R.H., A.J.S.J., C.T., A.M.R., J.P.S., C.A.W., L.Z., S.S.W.; Resources, C.J.G., M.S.M., L.W.M., S.S.W.; Data Curation, L.A.B, J.R.H., A.J.S.J., A.M.R., J.P.S., C.T., B.J.Z., L.Z., S.S.W.; Writing, A.M.R., J.P.S., B.J.Z., S.S.W.; Reviewing and Editing, All authors; Visualization, A.M.R., J.P.S., B.J.Z., S.S.W.; Supervision, C.J.G., S.S.W.; Project Administration, C.J.G., M.C.K., S.S.W.; Funding Acquisition, C.J.G., S.S.W.. *Funding support*: This work was supported by the National Heart, Lung, and Blood Institute (R01HL158647 to S.S.W., R01HL167969 to C.J.G. and L.W.M., and R01HL180692 to I.K.) and the National Institute of Arthritis and Musculoskeletal and Skin Diseases (T32AR065971 to A.J.S.J. and C.T.) of the National Institutes of Health. Additional support was provided by the Indiana University School of Medicine (to J.P.S.), and Purdue University College of Engineering and Indiana University School of Medicine Pilot Grant Initiation for Engineering in Medicine Pilot Project (to S.S.W., C.J.G. and L.W.M.), and the Riley Children’s Foundation (to S.S.W., C.J.G., M.C.K. and L.W.M.).

## Acknowledgements

The authors thank the members of the Indiana University School of Medicine Histology Lab Service Core and Center for Diabetes and Metabolic Diseases Translational Core for their outstanding technical support.

