## Supplementary Figures for "Empagliflozin preserves cardiac function and modulates metabolism in a mouse model of Duchenne muscular dystrophy"

**Supplementary Figure 1: Cardiac mass and volume metrics demonstrate consistent temporal changes across groups.** (A) *Ex vivo* heart mass measured after euthanasia. (B) Left ventricular (LV) mass derived from 4D ultrasound segmentation. (C) LV end-diastolic volume. (D) LV peak systolic volume. Data are presented as mean ± SD. N = 9–10 per group. Multiple group comparisons were analyzed using ordinary one-way ANOVA with Tukey’s post hoc test and longitudinal comparisons were analyzed using two-way repeated-measures ANOVA with Tukey’s post hoc test. Statistical significance: ^, *P* < 0.05, *mdx* vs. WT; *, *P* < 0.05, WT vs. *mdx* and *mdx* + EMPA; #, *P* < 0.05, all groups significantly different from one another.

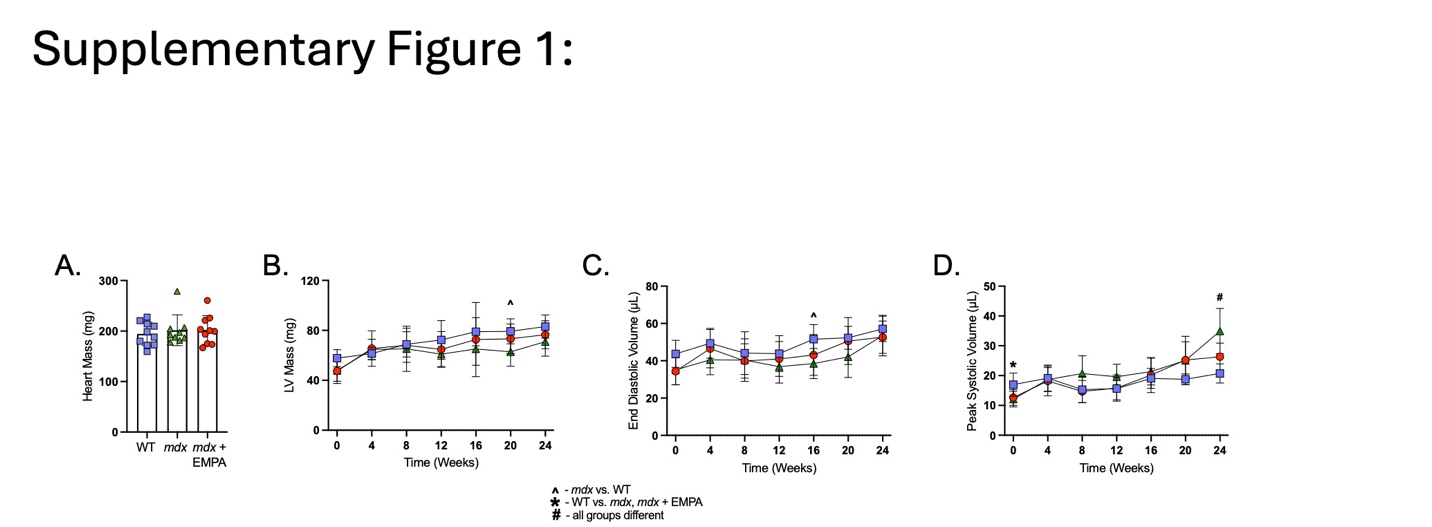

**Supplementary Figure 2:** **Group differences in longitudinal strain reveal delayed divergence in *mdx* mice with empagliflozin treatment.** Group-averaged heatmaps of differences in LV longitudinal strain between WT + EMPA, *mdx*, and *mdx* + EMPA groups across all time points. Regional strain (y-axis: free wall, anterior, septum, posterior, and free wall) is displayed across one cardiac cycle (x-axis). Minimal differences are observed between WT + EMPA and *mdx* + EMPA until week 16, whereas greater divergence is evident between WT + EMPA and *mdx* groups.

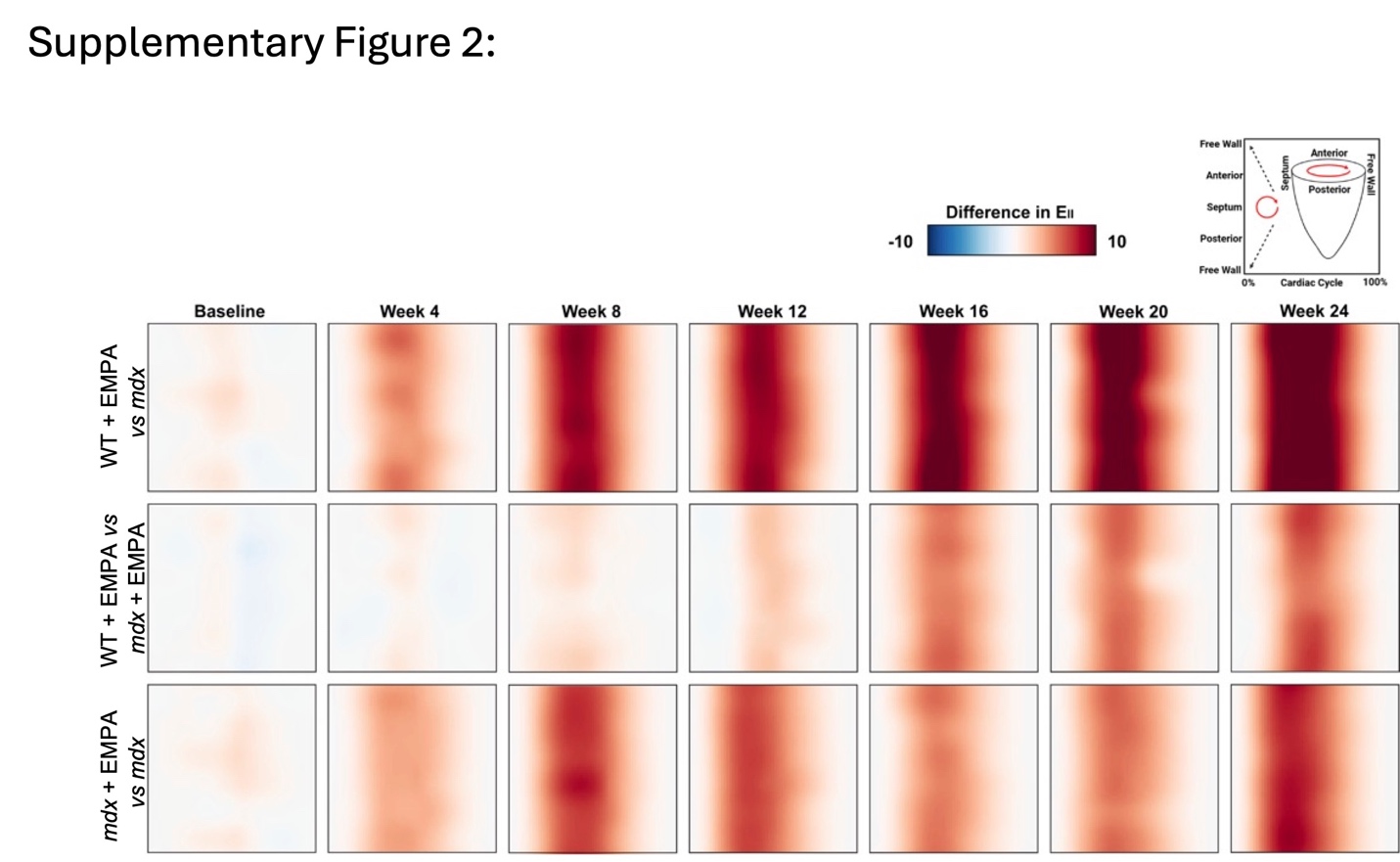

**Supplementary Figure 3: Regional longitudinal strain metrics are consistent across regions and agree with global trends.** The left ventricle (LV) was segmented into 6 regions (clockwise): anterior wall (A), anterior free wall (AFW), posterior free wall (PFW), posterior wall (P), posterior septum (PS), anterior septum (AS). (A) Regional peak longitudinal strain. (B) Regional systolic longitudinal strain rate. (C) Regional early diastolic longitudinal strain rate. (D) Regional late diastolic longitudinal strain rate. Data are presented as mean ± SD. N = 9–10 per group. Longitudinal comparisons were analyzed using two-way repeated-measures ANOVA with Tukey’s post hoc test. Statistical significance: x, *P* < 0.05, *mdx* vs. WT and *mdx* + EMPA; ^, *P* < 0.05, *mdx* vs. WT; #, *P* < 0.05, all groups significantly different from one another.

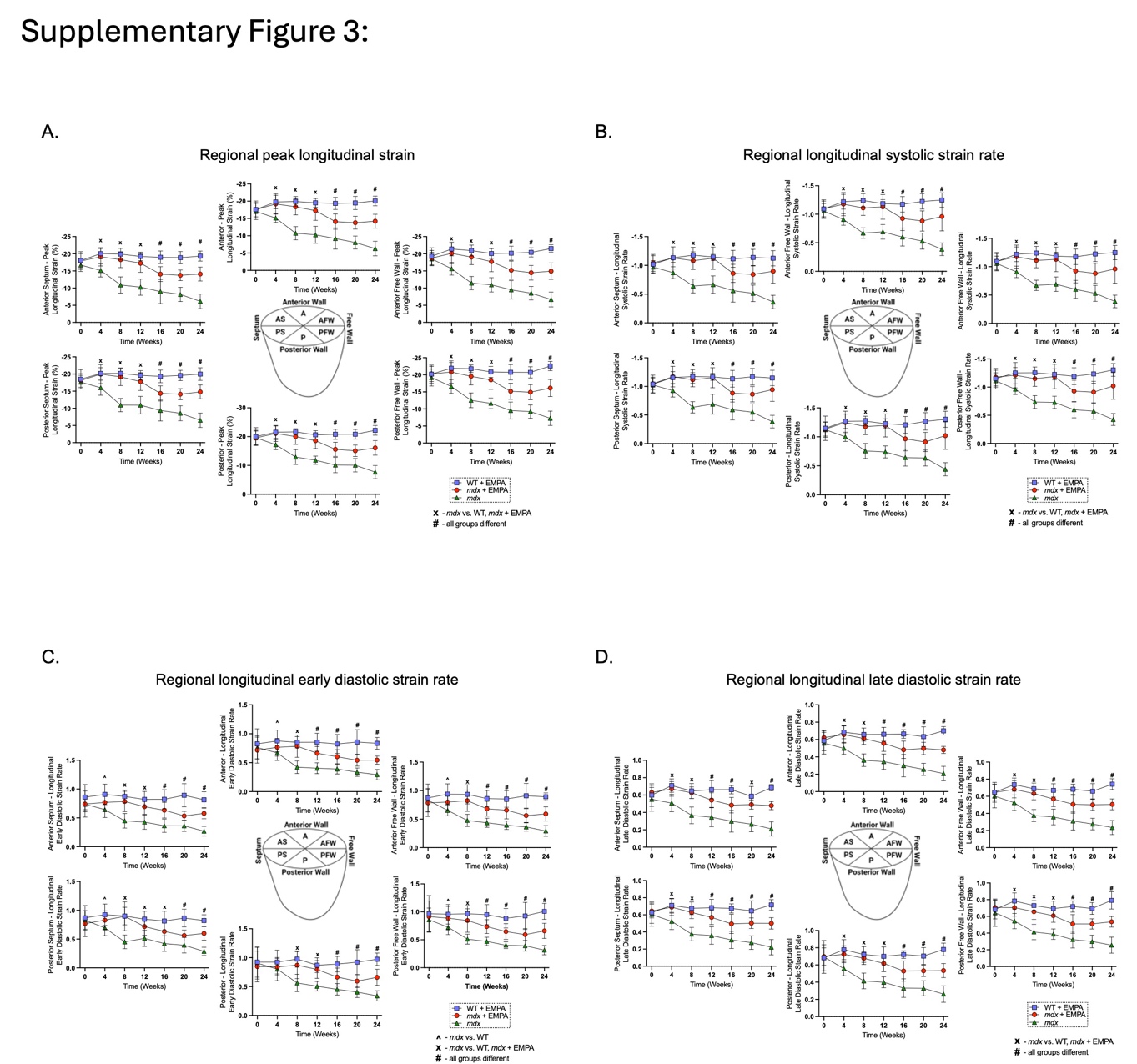

**Supplementary Figure 4: Circumferential and surface area strain metrics follow global longitudinal strain metrics.** (A) Global circumferential peak strain, systolic strain rate, early diastolic strain rate, and late diastolic strain rate over 24 weeks. (B) Global surface area peak strain, systolic strain rate, early diastolic strain rate, and late diastolic strain rate over 24 weeks. Data are presented as mean ± SD. N = 9–10 per group. Longitudinal comparisons were analyzed using two-way repeated-measures ANOVA with Tukey’s post hoc test. Statistical significance: x, *P* < 0.05, *mdx* vs. WT and *mdx* + EMPA; ^, *P* < 0.05, *mdx* vs. WT; *, *P* < 0.05, WT vs. *mdx* and *mdx* + EMPA; #, *P* < 0.05, all groups significantly different from one another.

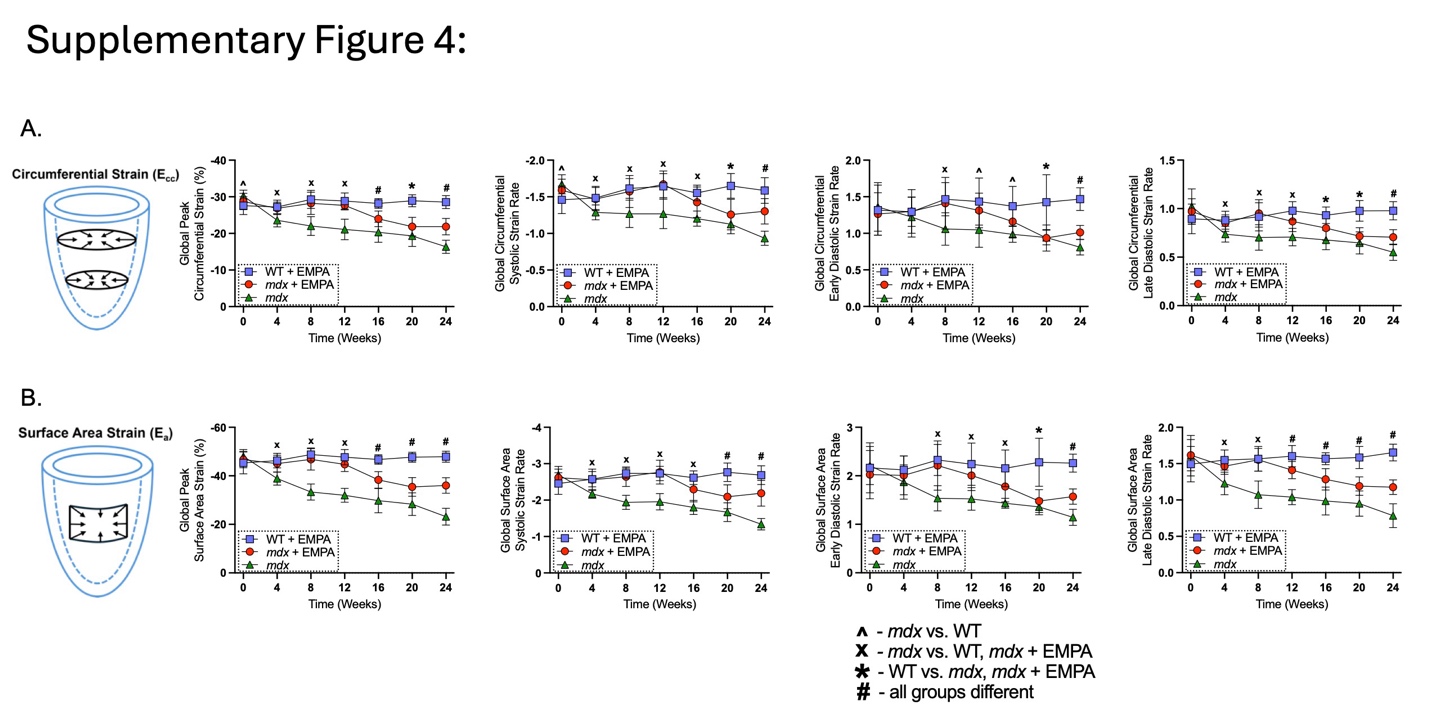

**Supplementary Figure 5: Regional circumferential strain metrics demonstrate homogeneous pattern of preservation from base to apex.** Regional circumferential peak strain, systolic strain rate, early diastolic strain rate, and late diastolic strain rate metrics at the basal, mid-ventricular, and apical levels of the LV over 24 weeks. Data are presented as mean ± SD. N = 9–10 per group. Longitudinal comparisons were analyzed using two-way repeated-measures ANOVA with Tukey’s post hoc test. Statistical significance: x, *P* < 0.05, *mdx* vs. WT and *mdx* + EMPA; ^, *P* < 0.05, *mdx* vs. WT; *, *P* < 0.05, WT vs. *mdx* and *mdx* + EMPA; #, *P* < 0.05, all groups significantly different from one another.

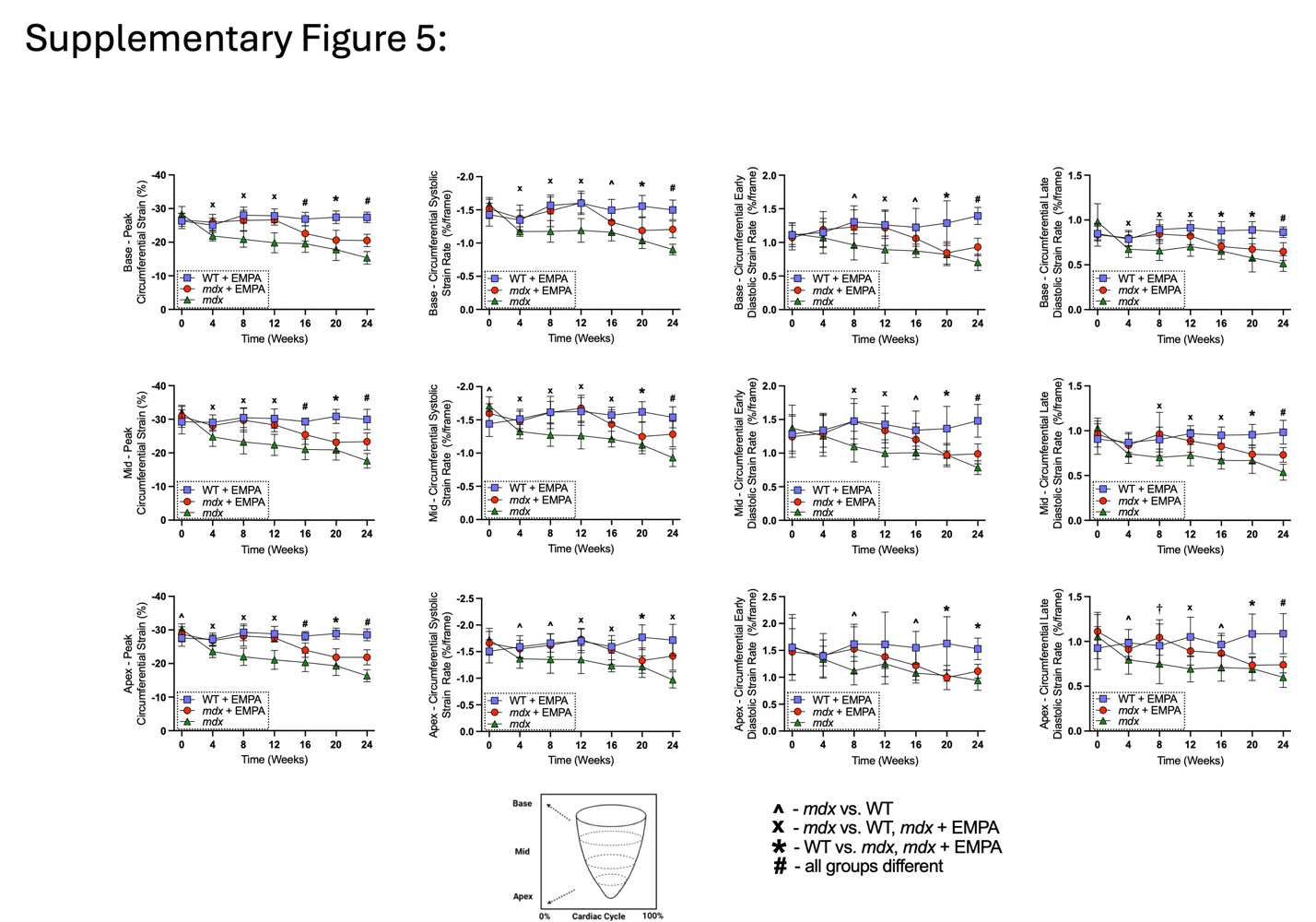

**Supplementary Table 1:** qRTPCR and QPCR Primers.

| ***Gene*** | **Forward Primer** | **Reverse Primer** |
| --- | --- | --- |
| *Ap3d1* | 5’-GATCATCAAGCTGTTCGGTGC-3’ | 5’-GAGATGAGCACAGCAATTACGG-3’ |
| *Bdh1* | 5’-AGGCTGTGACTCTGGATTTGGG-3’ | 5’-CTGGATGGTTCTCAGTCGGTCA-3’ |
| *Col1a1* | 5’-TGTGTGCGATGACGTGCAAT-3’ | 5’-GGGTCCCTCGACTCCTACA-3’ |
| *Col3a1* | 5’-ATCCCATTTGGAGAATGTTGTGC-3’ | 5’-GGACATGATTCACAGATTCCAGG-3’ |
| *Cpt1b* | 5’-CCCCTCATGGTGAACAGCAA-3’ | 5’-GGTCCAGTTTGCGGCGATAC-3’ |
| *Fn1* | 5’-GCTCAGCAAATCGTGCAGC-3’ | 5’-CTAGGTAGGTCCGTTCCCACTG-3’ |
| *Havcr1* | 5’-CTGGAATGGCACTGTGACATCC-3’ | 5’-GCAGATGCCAACATAGAAGCCC-3’ |
| *Hk2* | 5’- GCCAGCCTCTCCTGATTTTAGTGT -3’ | 5’- GGGAACACAAAAGACCTCTTCTGG -3’ |
| *Lcn2* | 5’-GAGCTACAATGTGCAAGTGG-3’ | 5’-GATGATGTTGTCGTCCTTGAG-3’ |
| *Lrp2* | 5’-AGAATGTGGCAGTGGGAATTT-3’ | 5’-GGACAGCCAATTTCATCAGTGT-3’ |
| *Myh6* | 5’-GCCCAGTACCTCCGAAAGTC-3’ | 5’-GCCTTAACATACTCCTCCTTGTC-3’ |
| *Myh7* | 5’-ACTGTCAACACTAAGAGGGTCA-3’ | 5’-GATGATTTGATCTTCCAGGG-3’ |
| *Nppa* | 5’-GCTTCTTCCTCGTCTTGGC-3’ | 5’-GGGGCATGACCTCATCTTC-3’ |
| *Nppb* | 5’-CTTTATCTGTCACCGCTGGGAG-3’ | 5’-TGCTGCCTTGAGACCGAAG-3’ |
| *Oxct1* | 5’-CATAAGGGGTGTGTCTGCTACT-3’ | 5’-CACATAGCCCAAAACCACCAA-3’ |
| *Pdk4* | 5’-TGAACACTCCTTCGGTGCAG-3’ | 5’-GCGTGTCTACAAACTCTGACA-3’ |
| *Ppara* | 5’-ATGCCAGTACTGCCGTTTTC-3’ | 5’-GGCCTTGACCTTGTTCATGT-3’ |
| *Psmc4* | 5’-CGACCTGGAAGACTATGTGGC-3’ | 5’-AGCCAACATTCCACTCTCCTG-3’ |
| *Rps20* | 5’-GAAGTCGCTGGAGAAGGTTTG-3’ | 5’-TCTTGGTAGGCATGCGCAC-3’ |
| *Slc16a1* | 5’-TGGTGTTATTGGAGGTCTTGG-3’ | 5’-CTGATTAAGTGGAGCCAGGGT-3’ |
| *Slc2a4* | 5’-GGTGTGGTCAATACGGTCTTCAC-3’ | 5’-AGCAGAGCCACGGTCATCAAGA-3’ |
| *Slc5a2* | 5’-GTGTTGGCTTGTGGTCTATGT-3’ | 5’-ATGACCGCTGCCGATGTTG-3’ |
| *Ywhaz* | 5’-GCCTGCTCTCTTGCAAAAACAG-3’ | 5’-GGGTATCCGATGTCCACAATG-3’ |
| *16S rRNA* | 5’- CCGCAAGGGAAAGATGAAAGAC-3’ | 5’- TCGTTTGGTTTCGGGGTTTC-3’ |
| *10 kb mito fragment* | 5’-GCCAGCCTGACCCATAGCCATAATAT-3’ | 5’-GAGAGATTTTATGGGTGTAATGCGG-3’ |
| *117 bp mito fragment* | 5’-CCCAGCTACTACCATCATTCAAGT-3’ | 5’-GATGGTTTGGGAGATTGGTTGATGT-3’ |
